# Robust and fast multicolor Single Molecule Localization Microscopy using spectral separation and demixing

**DOI:** 10.1101/2023.01.23.525017

**Authors:** Karoline Friedl, Adrien Mau, Valentina Caorsi, Nicolas Bourg, Sandrine Lévêque-Fort, Christophe Leterrier

## Abstract

Single Molecule Localization Microscopy (SMLM) is a straightforward approach to reach sub-50 nm resolution using techniques such as Stochastic Optical Reconstruction Microscopy (STORM) or DNA-Point Accumulation for Imaging in Nanoscale Topography (PAINT), and to resolve the arrangement of cellular components in their native environment. However, SMLM acquisitions are slow, particularly for multicolor experiments where channels are usually acquired in sequence. In this work, we evaluate two approaches to speed-up multicolor SMLM using a module splitting the fluorescence emission toward two cameras: simultaneous 2-color PAINT (S2C-PAINT) that images spectrally-separated red and far-red imager strands on each camera, and spectral demixing STORM (SD-STORM) that uses spectrally-close far-red fluorophores imaged on both cameras before assigning each localization to a channel by demixing. For each approach, we carefully evaluate the crosstalk between channels using three types of samples: DNA origami nanorulers of different sizes, single-target labeled cells, or cells labeled for multiple targets. We then devise experiments to assess how crosstalk can potentially affect the detection of biologically-relevant subdiffraction patterns. Finally, we show how these approaches can be combined with astigmatism to obtain three-dimensional data, and how SD-STORM can be extended three-color imaging, making spectral separation and demixing attractive options for robust and versatile multicolor SMLM investigations.

## Introduction

The advent of super-resolution microscopy has allowed to bypass the diffraction barrier in fluorescence imaging and study the fine details of the cellular architecture (Vangindertael et al., 2018; Jacquemet et al., 2020). Among the various techniques available to biologists, Single Molecule Localization Microscopy (SMLM) requires relatively simple equipment and can reach a lateral resolution below 50 nm (Sauer and Heilemann, 2017; Lelek et al., 2021; Liu et al., 2022). Its principle is to temporally decompose a fluorescent staining into sparsely blinking fluorophores, allowing their localization through single molecule detection and fitting on the successive frames of a continuous acquisition, the final image being generated by plotting all the fitted coordinates of the detected fluorophores (also called localizations) (Deschout et al., 2014). SMLM was first invented in the form of Stochastic Optical Reconstruction Microscopy (STORM), where organic fluorophores sparse blinking is induced by high-power laser illumination and a reducing buffer (Rust et al., 2006; Heilemann et al., 2008), and as Photoactivate Localization Microscopy (PALM) where the blinking of photoactivable fluorescent proteins is induced by low-power illumination (Betzig et al., 2006; Hess et al., 2006). A third SMLM approach was later developed where the blinking is generated from the transient interaction of short fluorescent DNA strands with their cognate docking strand conjugated to antibodies, a technique called DNA-Point Accumulation for Imaging in Nanoscale Topography (PAINT) (Jungmann et al., 2014).

The single-molecule nature of SMLM allows for its exquisite resolution, but this comes with two important consequences: it is an inherently slow technique, where fluorophores must be localized one by one; and distinguishing targets based on their labeling with different fluorophores is not as straightforward as with ensemble fluorescence techniques. Efforts to speed up STORM have focused on higher power lasers to accelerate blinking (Lin et al., 2015), but this can limit the total amount of localizations retrieved (Diekmann et al., 2020); deep-learning based algorithms were also developed that can work with higher blinking densities (Nehme et al., 2018; Speiser et al., 2021) or infer images from a subset of acquired frames (Ouyang et al., 2018). Multicolor STORM was first demonstrated with antibodies conjugated with different couples of activator and reporter fluorophores (Bates et al., 2007, 2012), while the search was ongoing to find good blinking probes outside of the far-red part of the spectrum for multicolor direct STORM (Dempsey et al., 2011; Lehmann et al., 2016; Wang et al., 2022). Alternatively, sequential acquisition schemes using the same fluorophore over cycles of quenching and restaining where developed (Tam et al., 2014; Valley et al., 2015; Yi et al., 2016). More complex approaches were also developed such as spectral STORM that distinguishes fluorophores based on their individual spectrum (Zhang et al., 2015, 2019; Dong et al., 2016; Yan et al., 2018), fluorophore distinction based on fluorescence lifetime (Thiele et al., 2020) or on differentially engineered point-spread functions (Shechtman et al., 2016; Wang et al., 2021; Eynde et al., 2022).

Among these approaches, a simple and elegant idea was to take advantage of the isolated nature of single molecule blinking events and determine their identity by splitting the emitted fluorescence in two paths using a dichroic mirror, identifying fluorophores based on the balance of intensities detected on each side. This “spectral demixing” strategy was developed soon after SMLM invention, using spectrally-close fluorophores illuminated by a single laser in different spectral bands or even fluorescent proteins (Schönle and Hell, 2007; Bossi et al., 2008; Testa et al., 2010; Baddeley et al., 2011; Gunewardene et al., 2011; Lampe et al., 2012). The optimal combination later converged toward 2- to 4-color imaging using far-red fluorophores such as DY634, Alexa Fluor 647 (AF647), CF647, DL650, CF660C or CF680 (Platonova et al., 2015; Winterflood et al., 2015; Lehmann et al., 2016; Favuzzi et al., 2017; Gorur et al., 2017; Zhang et al., 2020; Andronov et al., 2022; Siemons et al., 2022). A key feature of this spectral-demixing STORM (SD-STORM) approach is the simultaneous imaging of all fluorophores, providing sped-up acquisition at the cost of a higher blinking density to be managed by the processing algorithm, which should be able to pair blinking events on images from both paths to reliably assign them to each channel (Tadeus et al., 2015; Andronov et al., 2022; Li et al., 2022).

Multiplexing PAINT for multicolor imaging is more straightforward than STORM, as it is possible to devise orthogonal DNA sequences and image them successively, washing out the previous imager strand before a new round of imaging - a method called Exchange-PAINT (Jungmann et al., 2014; Agasti et al., 2017; Guo et al., 2019). However, this results in extremely long acquisition times, blinking in PAINT still being inherently slower than STORM, despite efforts to speed up the transient binding of imager strands (Schueder et al., 2019; Civitci et al., 2020; Strauss and Jungmann, 2020). A couple of recent attempts have been made to speed up PAINT through simultaneous multicolor acquisitions, either spectrally separated (Cheng et al., 2021; Chung et al., 2022) or using spectral demixing (Gimber et al., 2022).

We recently developed an SMLM module that couples large field-of-view illumination (Mau et al., 2021) to a 2-way detection path with a dichroic mirror splitting fluorescence toward two cameras. Here, we used this module to implement two approaches for fast 2-color SMLM via either spectral separation (red and far red fluorophores excited by two lasers) or spectral demixing (two far red fluorophores excited by a single laser), and we compared their performance. We developed innovative samples and procedures to assess the crosstalk between channels in both modalities, and evaluate its influence on biologically-relevant imaging experiments. Finally, we demonstrate that both simultaneous 2-color PAINT and SD-STORM are readily extensible to astigmatism-based 3D SMLM, whereas SD-STORM extension to three colors is straightforward.

## Results

### Simultaneous two-color DNA-PAINT imaging

Our first strategy for simultaneous multicolor super-resolution imaging is based on the traditional principle of spectral separation in fluorescence microscopy. This is done using a pair of fluorophores that have distinct excitation and emission spectra: Cy3B or Atto565 (emitting in the red part of the spectrum) and Atto643 or Atto655 (emitting in the far-red), with their emission split by a dichroic towards two cameras. The dichroic mirror splits the emitted fluorescence at 662 nm, reflecting the red fluorophore emission to one camera and transmitting the far-red fluorophore emission to the other camera (Fig. S1). Fluorophore excitation occurs via continuous illumination with two laser sources at 532 nm and 640 nm (Fig. S1A). A STORM-based implementation of this 2-color imaging approach does not provide high-quality images, due to the difficulty of finding a STORM buffer optimal for both a red and a far-red fluorophore, and the lower blinking quality of red fluorophores including Cy3B and CF568 (Dempsey et al., 2011; Lehmann et al., 2016)(Fig. S2). By contrast, DNA-PAINT is a good candidate for 2-color imaging by spectral separation, as blinking from the red and far-red strands hybridization occurs with similar characteristics, and the blinking density can be independently adjusted by modifying each imager strand concentration (Jungmann et al., 2014; Schnitzbauer et al., 2017). Most multicolor DNA-PAINT studies have used sequential acquisition of targets by successive incubation with different imagers conjugated to the same fluorophore. However, it should be straightforward to increase the acquisition speed by simultaneously acquiring two channels using spectrally-separated fluorophore on distinct imagers, as demonstrated recently (Cheng et al., 2021; Chung et al., 2022).

We thus performed simultaneous 2-color acquisition by DNA-PAINT (S2C-PAINT) using imager strands conjugated with fluorophores emitting in the red channel (Cy3B or Atto 565) and far-red channel (Atto643 or Atto655) (Fig. 1A). To check if some crosstalk can occur for this 2-color approach and compare it rigorously to the spectral demixing strategy, we performed a ratiometric analysis (Baddeley et al., 2011; Lampe et al., 2012): blinking events were processed, resulting in localizations coordinates. Localizations appearing at the same time and within 500 nm of each other on each camera frame were paired, and their ratio of photons was calculated by dividing the number of photons in the transmitted pathway T (corresponding to the far-red channel) by the total number of photons found on both cameras R+T (R: red channel). When present only on the frame from the reflected pathway (red channel), a localization was assigned a ratio of 0; whereas if it is found only on the frame from the transmitted pathway, it was assigned a ratio of 1. The average photon ratio distribution from several S2C-PAINT acquisitions (microtubules and clathrin in COS cells, Fig. 1D) with the Cy3B/Atto643 imager pair is shown in Fig. 1A, lower right. Only ~3% of the total localizations are paired, showing that most blinking events only appear on one camera. Accordingly, we chose to assign localizations with ratios between 0 and 0.01 to the red channel, and between 0.99 and 1 for the far-red channel (Fig.1A, lower right).

**Figure 1:**
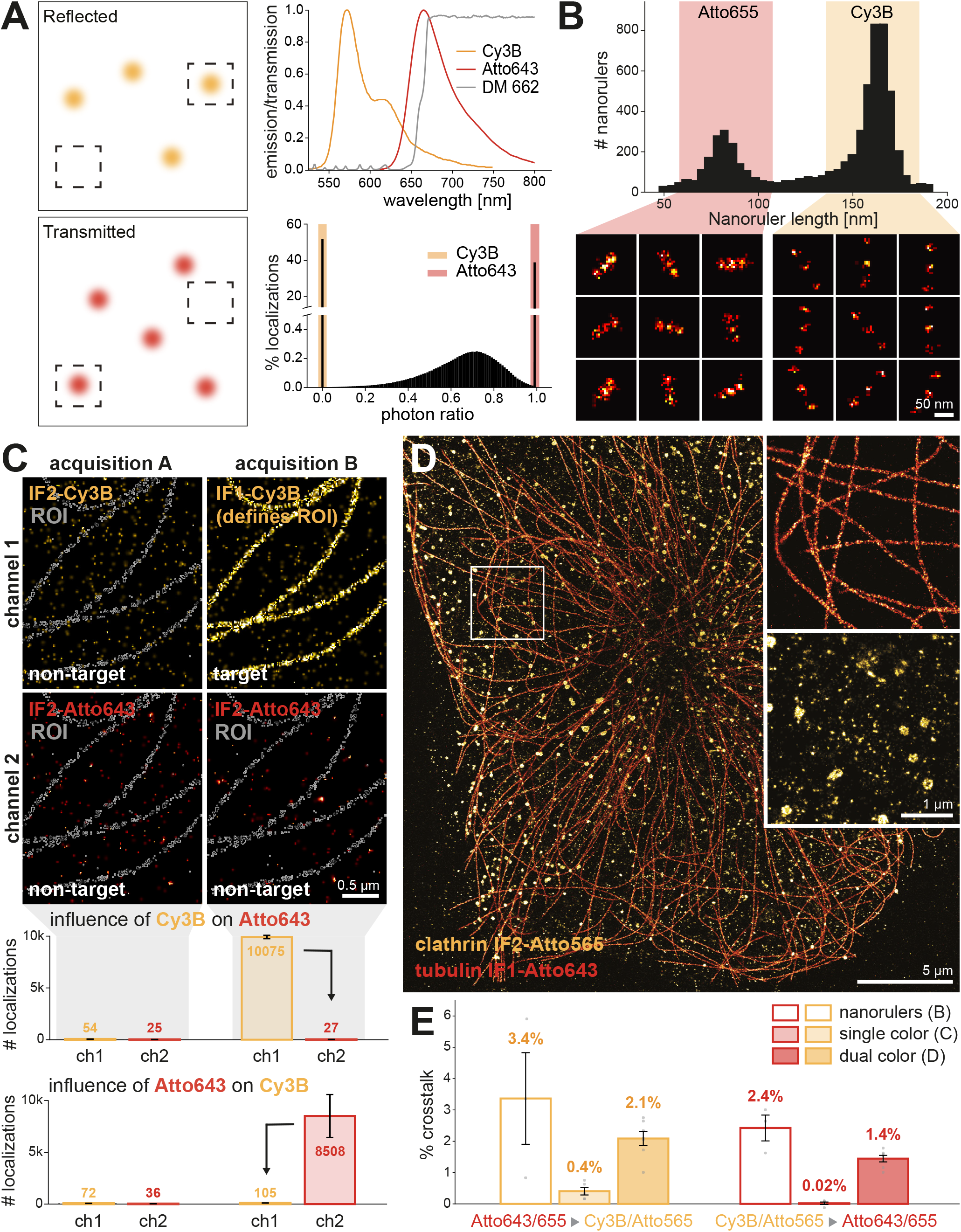
Simultaneous two-color DNA-PAINT (S2C-PAINT) and crosstalk evaluation. **A.** In S2C-PAINT, blinking events appear only on either one camera or the other (left panels) because of the large spectral separation between fluorophores (top right graph): fluorophores emitting in the red (here Cy3B, orange curve) and far-red (here Atto643, red curve) are separated by the 662 nm cut-off dichroic mirror (gray line). Ratiometric analysis confirms that most fluorophores appear only on one camera (bottom right graph): the range of ratio associated to Cy3B is chosen between 0.00 and 0.01; the range associated with Atto643 is between 0.99 and 1.0 (orange and red colored areas). **B.** Crosstalk measurement from nanorulers: simultaneous 2-color imaging of a slide combining two types of three-spots nanorulers (40-nm and 80-nm spacing) labeled with P1 and P3 PAINT docking strands respectively, with a mixture of I1-Atto655 and I3-Cy3B imager strands. Nanorulers were classified by total length (top distributions, peaks at 80 and 160 nm) to determine the crosstalk (see E and Figure S3). **C.** Crosstalk measurement from a single-target cellular sample: zoomed image of a COS cell labeled for microtubules with a secondary antibody carrying an F1 docking strand. Acquisition A (top, left column) uses a mixture of non-target (IF2) imager strands to measure the background signal in both channels. Acquisition B (top, right column) is done over the same field of view with a mixture of a target (IF1-Cy3B) and a non-target (IF2-Atto643) imager strand. A ROI enclosing the microtubule network (bottom row, white is a ROI overlay on the background images) is obtained from the target (IF1-Cy3B) image and used to measure the number of localizations for Acquisition A and B in both channels (middle graph). The crosstalk from Cy3B into Atto643 is based on the difference between the number of localizations in acquisition B, channel 2 (27, right red bar) and the number of localizations in acquisition A, ch2 (25, left red bar). Conversely, the crosstalk of Atto643 into Cy3B can be measured using a target IF2-Cy3B acquisition, evaluating the difference induced in ch1 between acquisition A and B (orange bars in bottom graph). See Fig. S4 for more details. **D.** Crosstalk measurement from a two-target cellular sample: COS cells labeled for tubulin (F1 docking strand) and clathrin (F2 docking strand), simultaneously imaged with IF1-Atto643 and IF2-Atto565. Insets show zoomed isolated channels. Crosstalk was calculated from exclusive ROIs containing only microtubules or only clathrin-coated pits excluding overlapping areas (see Fig. S5). **E.** Crosstalk values obtained from the three different approaches, grouped by crosstalk direction (far-red into red on the left, orange; red into far-red on the right, red): nanorulers (see B), single-target cellular sample (C), and two-target cellular sample (D). Points corresponds to individual images, error bars are SEM.

### Measurement of crosstalk on three different types of samples

We then tested several types of samples that allowed us to assess the performance and potential crosstalk of the S2C-PAINT approach on distinct types of samples: a mix of DNA origami nanorulers of different sizes (Lin et al., 2019; Scheckenbach et al., 2020)(Fig. 1B), cellular samples stained against a single-target (tubulin in COS cells, Fig. 1C), or against two targets (clathrin and tubulin in COS cells, Fig. 1D)(Jimenez et al., 2020). The custom nanoruler slide combines two types of 3-spot rulers: 80-nm long nanorulers with a 40 nm distance between the spots labeled with a P1 docking strand (“short” nanoruler), and 160-nm long nanorulers with an 80-nm distance between the spots labeled with a P3 docking strand (“long” nanoruler). The nanorulers were imaged simultaneously using I1-Atto655 (short) and I3-Cy3B (long) fluorescent imager strands (Fig. 1B). We devised these nanorulers to obtain structures identified solely from their shape (length and cluster spacing) without relying on channel information, providing an independent way of measuring crosstalk. Individual nanorulers were segmented by density-based clustering (DBSCAN) (Ester et al., 1996) and classified based on their total length (Fig. S3). Downstream of this shape-based classification, we then calculated the crosstalk between channels by measuring the number of localizations found in the non-labeled channel divided by the number of localizations in both the labeled and non-labeled channels. For example, we measured the crosstalk of the Atto655 channel into the Cy3B channel by measuring the number of localizations in the Cy3B channel (I3-Cy3B imager) divided by the number of localizations in both the Cy3B channel and the Atto655 channel (I1-Atto655 imager) on the short nanorulers (P1 docking strand). We found this crosstalk from the Atto655 channel into the Cy3B channel to be 3.37% (Fig. 1E, left), whereas the crosstalk from Cy3B channel into the Atto655 channel was found to be 2.42% (Fig. 1E, right).

A more classic way of assessing crosstalk is to use single-target labeling in cells, and to measure the signal induced in the non-target channel by the labeling in the target channel. This is however not applicable when using high-performance SMLM algorithms that detect blinking events based on a relative threshold, as the detection threshold will be lowered in the non-target channel, resulting in the detection of spurious localization events and a significantly overestimated crosstalk. This is especially true when using DNA-PAINT, as the local variation of the fluorescent imager background will provide fitting candidates to an adaptive algorithm in the absence of true blinking events. To assess crosstalk from single-target samples using the same algorithm as for other samples, we devised a method based on repeated imaging of the same field of view with different labels present, a unique possibility offered by DNA-PAINT. We thus measured the change in localization numbers for a non-target (“empty”) channel that is induced by having an imager strand present in the “target” channel: for example, we observed if more localizations would be detected in the “empty” red channel (devoid of imager), whether an Atto643 imager strand was present and interacted with its cognate docking strand on a cellular labeling in the far-red channel. We prepared COS cells stained for microtubules with an anti-tubulin antibody and a secondary antibody conjugated to an F1 docking sequence, then acquired two sequences. The first sequence was acquired using a mixture of IF2-Cy3B and IF2-Atto643 imager strands that both do not bind to the tubulin staining, measuring the number of localizations from the DNA-PAINT background in both channels (Fig. 1C, left column, “acquisition A”). To estimate the crosstalk from the Cy3B channel into the Atto643 channel, we then acquired a second sequence on the same field of view, this time with a mixture of the IF1-Cy3B and IF2-Atto643 imager strands (Fig. 1C, right column, “acquisition B”). In this second acquisition sequence, the microtubule staining is revealed in the Cy3B channel, whereas the Atto643 channel contains background plus the crosstalk from the Cy3B channel staining. We segmented the microtubule staining in the Cy3B channel (Fig. 1C, upper right panel), and used the resulting region of interest (ROI) to measure the number of localizations in the Atto643 channel from each of the two acquired sequences (Fig. 1C, white outlines in the lower row images and corresponding graph). The crosstalk was calculated as the difference in the number of localizations within this ROI in the Atto643 channel when the Cy3 channel contains a target (acquisition B) and non-target (acquisition A) imager strand, divided by the background-corrected number of localizations in both channels (see Fig. S4 and *Methods*). Conversely, we acquired another pair of sequences to measure the crosstalk from the Atto643 channel into the Cy3B channel on F1-labeled microtubules. For this, the same acquisition A was performed with IF2-Cy3B and IF2-Atto643 imager strands, while acquisition B was performed with the IF2-Cy3B and IF1-Atto643 imager strands (Fig. 1C, bottom graph). With this new approach, we found very low values of crosstalk with S2C-PAINT: the Atto643 channel crosstalk into Cy3B is 0.41%, while the Cy3B channel crosstalk into the Atto643 channel is 0.02% (Fig. 1E).

Finally, we aimed at directly measuring the crosstalk from simultaneously acquired 2-color DNA-PAINT images, as this would be the most straightforward approach. We prepared COS cells labeled for clathrin-coated pits (anti-clathrin primary antibody and secondary antibody conjugated with an F2 docking sequence) and for microtubules (anti-tubulin primary antibody and secondary antibody conjugated with an F1 docking strand). The sample was imaged with a mixture of IF2-Atto565 and IF1-Atto643, and the two channels were obtained after demixing as detailed above (ratio range 0.0-0.01 for the Atto565 channel, 0.99-1.0 for the Atto643 channel). The reconstructed images show no visible crosstalk between channels (Fig. 1D) and a good structural quality for both targets, similar to what we can obtain with alternatively-acquired PAINT channels (Jimenez Methods 2020). To measure the crosstalk from this data, we generated ROIs that contained localizations from a single target, excluding regions where pits and microtubules overlap. To do this, we generated two ROIs by thresholding the microtubule network and the clathrin-coated pits, then derived exclusive-target ROIs by excluding overlapping areas using ROI subtraction (Fig. S5). We calculated the crosstalk in each exclusive-target ROI as the number of localizations within the exclusive-target ROI on the non-target image (example: microtubule-only ROI applied on the clathrin-coated pits image), divided by the sum of the localizations on the target (microtubules) and non-target (clathrin) images within the same ROI (example: sum of the localizations from the microtubules and clathrin-coated pits images within the microtubule-only ROI). We obtained a crosstalk value of 2.09% for the Atto643 channel crosstalk into the Cy3B channel, and of 1.44% for Cy3B into Atto643 (Fig. 1E).

A comprehensive overview of all acquisition parameters we used can be found in Table S1. Overall, the three different approaches yielded different crosstalk values for S2C-PAINT. We found higher crosstalk values from the nanorulers and two-target samples, and lower values from the background-corrected single-target samples. Nevertheless, the values are all quite low (below 4%), as expected from separating fluorophores with a large difference in emission peaks, and all three methods were consistent in detecting more crosstalk from the higher-wavelength channel (far-red Atto643/Atto655) into the lower-wavelength channel (red Cy3B/Atto565).

### Simultaneous 2-channel imaging using spectral demixing of two far-red fluorophores

The second strategy we implemented on our setup is to spectrally demix two far-red fluorophores: excited by a single laser illumination wavelength, blinking events from both fluorophores appear on each camera with a different photon ratio (Fölling et al., 2008; Testa et al., 2010; Baddeley et al., 2011; Lampe et al., 2012). In our implementation, the sample is illuminated with a single laser line at 640 nm and fluorophore emission is divided using a 700-nm long-pass dichroic mirror into a reflected and transmitted pathway (Fig. 2A and Fig. S1B), with most blinking events observed on the two cameras (Fig. 2A, left). Ratiometric analysis was performed as for S2C-PAINT imaging (see above): blinking events were detected and localized on the images from both cameras, with the number of photons measured for each blinking event. Localizations were then paired between corresponding camera images, and the photon ratio calculated using T/(R+T) (see *Methods*). The average photon ratio distribution from several SD-STORM acquisitions (microtubules and clathrin in COS cells, Fig. 2D) using AF647 and CF680 fluorophores showed a photon ratio peak at ~0.25 for AF647 (more photons on the reflected pathway camera) and ~0.55 for CF680 (equal repartition on both cameras, Fig. 2A, bottom right). To retain an optimal number of localizations from both channels and ensure structural quality of the images, we assigned the ratios between 0.01 and 0.38 to AF647 and those between 0.42 to 0.99 to CF680 (Fig. 2A, bottom right), excluding 1.4% localizations outside of the selected ranges, and the localizations only detected on a single camera as background (photon ratio of 0 or 1, which represent 33% and 10% of the total number of localizations on average in 2-color acquisitions).

**Figure 2:**
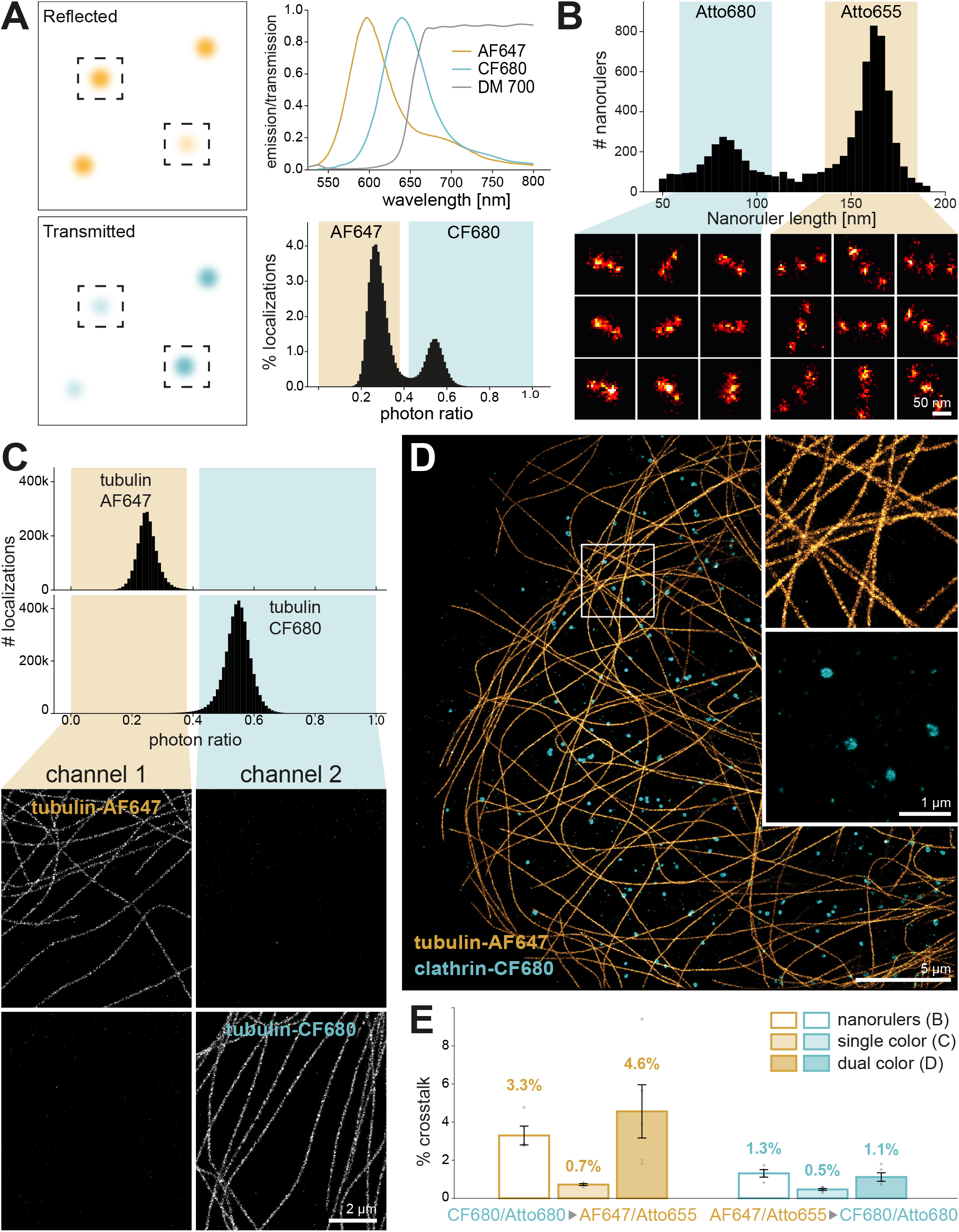
Spectral demixing STORM (SD-STORM) and crosstalk evaluation. **A.** In SD-STORM, blinking events from two far-red fluorophores (AF647 and CF680) appear on both cameras with different intensity ratios (left panels) because of the splitting through their close emission spectra by a 700 nm dichroic mirror (top right graph). Ratiometric analysis reveals characteristic populations of ratios for blinking events from each fluorophore (bottom right graph): the ratio population between 0.01 and 0.38 is associated to AF647 (yellow area), the ratio population between 0.42 and 0.99 is associated to CF680 (blue area). **B.** Crosstalk measurement from nanorulers: spectral demixing PAINT imaging of a slide combining two types of three-spots nanorulers (40-nm and 80- nm spacing) labeled with P1 and P3 PAINT docking strands respectively, with a mixture of I1-Atto680 and I3-655 imager strands. Channels were generated after demixing with the photon ratio ranges as specified in A. Nanorulers were classified by total length (top distribution, peaks 80 and 160 nm) to determine the crosstalk (see E and Figure S3). **C.** Crosstalk measurement from a single-target cellular sample: COS cells labeled for microtubules with a secondary antibody conjugated to either AF647 or CF680. Ratiometric analysis of the localizations (top) shows distinct populations for each fluorophore. After demixing (yellow and blue areas), images were reconstructed (bottom panels. ROIs were drawn around the microtubules with an intensity threshold and used on both channels to calculate the crosstalk (see E). **D.** Crosstalk measurement from a two-target cellular sample: COS cells labeled for tubulin with AF647 and for clathrin with CF680. Demixing was performed using the photon ratio ranges specified in A. Insets show zoomed isolated channels. Crosstalk was calculated from exclusive ROIs containing only microtubules or only clathrin-coated pits excluding overlapping areas (see Fig. S5). **E.** Crosstalk values obtained from the three different approaches, grouped by crosstalk direction (far-red into red on the left, orange; red into far-red on the right, red): nanorulers (see B), single-target cellular sample (C), and two-target cellular sample (D). Points corresponds to individual images, error bars are SEM.

To assess the crosstalk between the two channels in this spectral-demixing STORM (SD-STORM) approach, we used the same three types of samples as for S2C-PAINT: nanorulers, sample labeled for a single target, or sample labeled for two targets. For the nanorulers experiment, we used the same short and long nanorulers as with S2C-PAINT with three spots spaced by 40 and 80 nm, respectively (Fig. 2B). As these are PAINT samples conjugated to docking strands rather than organic fluorophores, we used an SD-PAINT imaging approach with two far-red imager strands: I1-Atto680 for the short nanoruler (40 nm spaced, P1 docking strand) and I3-Atto655 for the long nanoruler (80 nm spaced, P3 docking strand). After segmentation, the nanorulers were classified based on their total length, independently of their content in each channel (Fig. 2B, top and Fig. S3). The two fluorophores have spectra that are close to the ones of AF647 and CF680, allowing to use the same photon ratio range for demixing (0.01-0.38 for Atto655, 0.42-0.99 for Atto680) and crosstalk measurement. This resulted in a crosstalk of 3.30% crosstalk from Atto680 to Atto655, and 1.31% from Atto655 to Atto680 (Fig. 2E).

In 2-color SD-PAINT and SD-STORM, fluorophores appear on both camera frames, hence there is no issue with adaptive SMLM algorithms picking up background in an “empty” channel such as in S2C-PAINT. We thus could use the classical approach of imaging COS cells stained for microtubules using primary antibodies against tubulin and secondary antibodies conjugated to either AF647 or CF680 (Fig. 2C). Photon ratios of single-target samples show a single peak around the expected ratio values (~0.25 for AF647 and ~0.55 for CF680), allowing demixing using the previously determined ratio ranges (0.01-0.38 for AF647, 0.42-0.99 for CF680). To measure the crosstalk, we simply segmented microtubules on the image from the target channel, counted the localizations within this ROI in both channels (Fig. 2C, bottom panels), and expressed the crosstalk as the ratio of the localizations number in the non-target channel over the sum of the localization numbers in the non-target and target channels. This resulted in a crosstalk of 0.72% from the CF680 channel into the AF647 channel, and of 0.46% from the AF647 channel into the CF680 channel (Fig. 2E). Finally, we directly measured the crosstalk from 2-color SD-STORM acquisition of samples labeled for microtubules (AF647-conjugated secondary antibodies) and clathrin (CF680-conjugated secondary antibody). The merged reconstructed images (Fig. 2D) show a clean separation between the labeled structures, the zooms on the right side show the channels of microtubules and clathrin, separately. Like for S2C-PAINT, we calculated the crosstalk after generating single-target ROIs that excluded areas of overlap between microtubules and clathrin-coated pits (Fig. S5). This resulted in a crosstalk of 4.56% from the CF680 channel to the AF647 channel, and 1.12% from the AF647 channel into the CF680 channel (Fig. 2E).

Overall, each method for estimating crosstalk gives different results, highlighting their complementarity and the need to properly define how the crosstalk is measured. In SDPAINT (for the nanorulers) and SD-STORM (for cellular samples), the preferential crosstalk direction is the same as in S2C-PAINT: we found the crosstalk from the higher-wavelength channel (Atto680/CF680) into the lower-wavelength channel (Atto655/AF647) to be slightly higher than in the other direction (lower-wavelength into higher wavelength). Overall, crosstalk values are below 5%, with slightly lower performance in simultaneous 2-color acquisitions between S2C-PAINT (2.1% – 1.4% for each direction) and SD-STORM (4.6% – 1.1%), consistent with the closer proximity of fluorophores in SD-STORM.

### Effect of crosstalk on the detection of biologically-relevant patterns

Measurement of crosstalk on specifically designed samples is important to benchmark the performance of multi-color SMLM strategies. However, we wanted to devise an experiment that could assess how crosstalk can perturb the visualization of biological structures, depending on the relative abundance of each target. For this, we turned to the membrane-associated periodic scaffold along the axon of neurons that is made of rings of adducin-associated actin spaced every 190 nm by a layer of spectrin tetramers, requiring super-resolution microscopy to be visible (Xu et al., 2013; Leterrier, 2021). When hippocampal neurons in culture are labeled for adducin and the center of the spectrin tetramer, this results in 190-nm periodic patterns of spectrin and adducin in antiphase, with spectrin bands and adducin bands alternating along the axon (Xu et al., 2013; Cabriel et al., 2019). We reasoned that crosstalk from one target (for example adducin) into the other (spectrin) would directly perturb the measured periodicity of the latter (spectrin), as it would result in the appearance of spurious localization in-between the periodic bands. Furthermore, the labeling abundance of the spectrin and adducin being roughly similar in standard labeling conditions, we should be able to modulate the crosstalk by varying the labeling of one target (changing imager strand concentration in PAINT of antibodies concentrations in STORM) and examine how this modulation affects the periodicity of the other target.

We thus labeled rat hippocampal neurons in culture for the carboxyterminus of ß2-spectrin and for adducin, and first imaged the resulting periodic patterns by S2C-PAINT: ß2-spectrin revealed with a F1-conjugated secondary antibody and a IF1-Cy3B imager, adducin revealed with a F2-conjugated secondary antibody and a IF2-Atto643 imager (Fig. 3). In the experiment shown in Fig. 3A-C, we repeatedly imaged the same field of view by S2C-PAINT (50,000 frames for each acquisition), keeping the ß2-spectrin labeling constant with the IF1-Cy3B imager at its reference level (100% i.e. 500 pM, Fig. 3A, top row), and varying the abundance of the adducin labeling by using a rising concentration of IF2-Atto643 imager (1%-6.25 pM; 10%-62.5 pM; 100%-625 pM, Fig. 3A, middle row). We measured the periodicity of each labeling by calculating the autocorrelation of intensity profiles along segments of axons (Fig. 3B), and measuring the amplitude of the first peak at 190 nm (Zhong et al., 2014)(see *Methods*) (Fig. 3C). Modulating the concentration of the IF2-Atto643 imager results in a gradual appearance of periodicity in the adducin channel, with the amplitude rising from 0.17 at 1% to 0.53 and 0.57 at 10% and 100%, respectively (Fig. 3B-C bottom, blue). Even at the maximum concentration, the adducin staining does not perturb the spectrin periodicity with an amplitude staying high at 0.6-0.7 (Fig 3B-C top, yellow). In the reverse experiment, we varied the ß2-spectrin labeling with a rising concentration IF1-Cy3B imager at its reference level (1%-5 pM, 10%-50 pM, 100%-500 pM, Fig. 3D, top row), and kept the adducin labeling constant (IF2-Atto643 imager concentration at 100%-625 pM, Fig. 3D, middle row). Note that in this case we were not able to relocate the field of view of the first condition, so the image is different (Fig. 3D). The periodicity of the spectrin labeling was already apparent at 1% labeling, and at 100% labeling, spectrin did not affect the periodicity of the adducin pattern (Fig. 3E). In fact, the amplitude of the autocorrelation for adducin was lower (0.14) at 1% spectrin labeling, while it is unaffected by a switch between 10% and 100% spectrin labeling (0.28 and 0.30, respectively, Fig. 3F). Overall, this shows that S2C-PAINT allows for the detection of biologically-relevant patterns at the nanoscale, being robust to variations in the abundance of labeled proteins.

**Figure 3:**
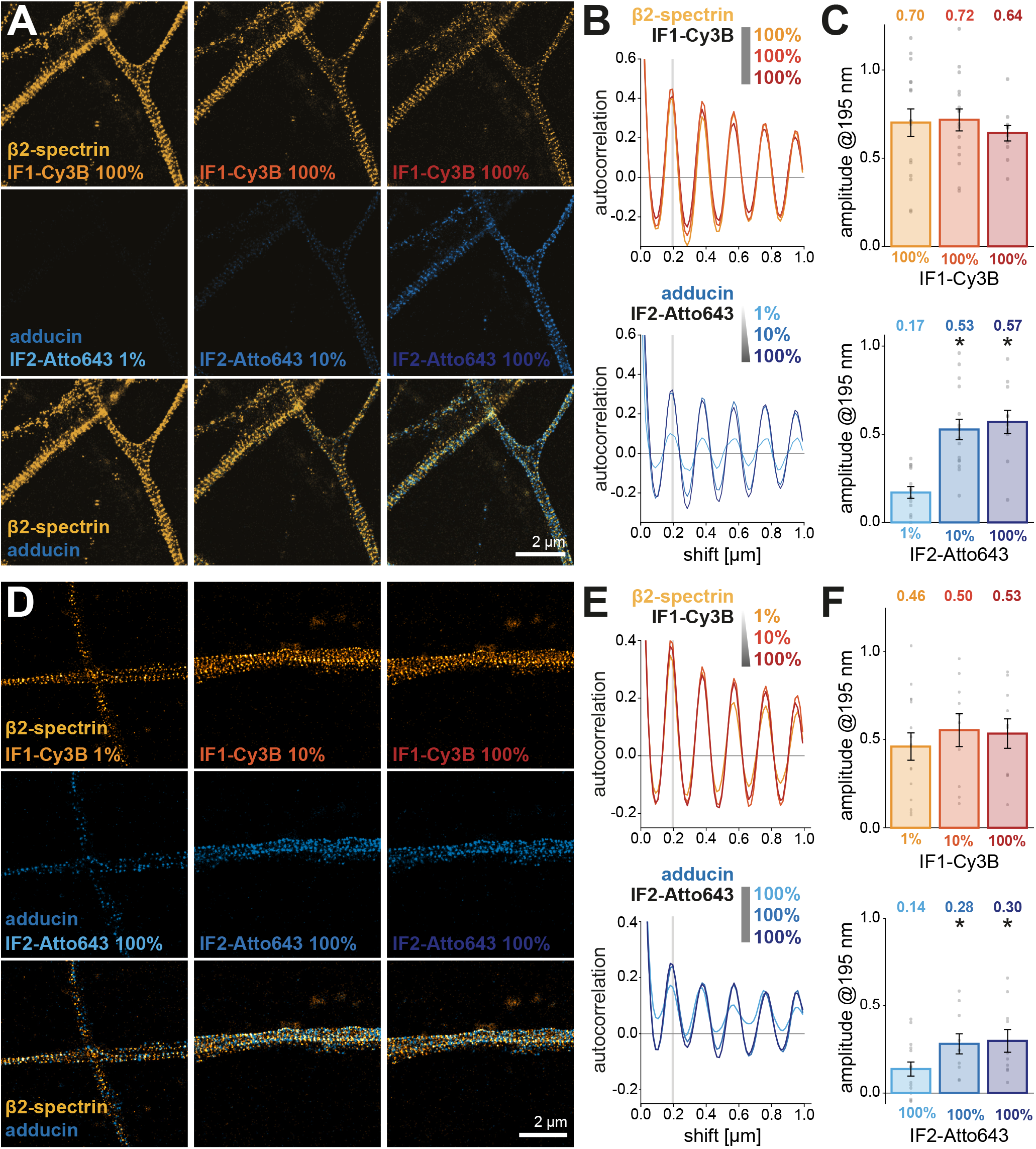
Effect of crosstalk on the detection of biologically-relevant patterns by S2C-PAINT. **A.** Reconstructed images from axons of hippocampal neurons stained for β2-spectrin (orange, top row) and adducin (blue, middle row), imaged by S2C-PAINT using IF1-Cy3B and IF2-Atto643, respectively. On the same field of view, the imager concentration of IF1-Cy3B (ß2-spectrin) was kept constant, while the concentration of IF2-Atto643 (adducin) was increased successively from 1% to 10% and 100% (columns) of the reference concentration (optimal imager concentration determined beforehand). **B.** Autocorrelation curves from 1-μm long intensity profiles along axons for the constant ß2-spectrin-IF1-Cy3B (orange, top) and varying adducin-IF2-Atto643 (bottom, blue) channels for each imager concentration conditions. **C.** Amplitude of the autocorrelation first peak (base-subtracted height at 195 nm shift) for each imager concentration value. Dots are individual axonal segments, bars are SEM. Stars show a significant difference with the 100%-1% condition (first bar) by post-hoc Tukey-Kramer test, p<0.05. **D.** Converse experiment where the ß2-spectrin imaging was done under varying IF1-Cy3B imager concentration (orange, top row) while adducin imaging was done at constant 100% IF2-Atto643 imager concentration (from 1% to 100%, blue, middle row). **E.** Autocorrelation curves from 1-μm long intensity profiles along axons for the varying ß2-spectrin-IF1-Cy3B (orange, top) and constant adducin-IF2-Atto643 (bottom, blue) channels for each imager concentration conditions. **F.** Amplitude of the auto-correlation first peak (base-subtracted height at 195 nm shift) for each imager concentration value. Dots are individual axonal segments, bars are SEM. Stars show a significant difference with the 100%-1% condition (first bar) by post-hoc Tukey-Kramer test, p<0.05.

We next performed similar experiments using SD-STORM, with adducin and ß2-spectrin labeled by secondary antibodies conjugated with AF647 and CF680, respectively. In SD-STORM, it is not possible to image the same field of view several times nor to modulate the density of labeling between acquisitions. We thus imaged sister coverslips immunolabeled with a mix of unconjugated and fluorophore-conjugated secondary antibodies containing 3.3%, 10% or 100% of conjugated antibody. We first kept the adducin-AF647 labeling constant at 100% and varied the ß2-spectrin-CF680 labeling from 3.3% to 100% (Fig. 4A-C). ß2-spectrin periodicity became detectable at 10% labeling (Fig. 4C bottom, blue), and the adducin labeling periodicity was unaffected by crosstalk from the ß2-spectrin labeling, with autocorrelation amplitudes staying constant between 0.21 and 0.28 (Fig. 4C top, yellow). In the reverse experiment, we kept the ß2-spectrin-CF680 labeling constant at 100% and varied the adducin-AF647 labeling from 3.3% to 100% (Fig. 4D-F). Here again, we detected a rise in the periodicity of the pattern for adducin between 3.3% and 100% (Fig. 4E top, yellow) and notably, we saw a drop in ß2-spectrin periodicity when increasing the adducin labeling density, from 0.30 at 1% to 0.25 at 10% and 0.16 at 100% adducin labeling (Fig. 4E bottom, blue). This is surprising because we consistently detected more crosstalk from the CF680 channel into the AF647 channel in our measurements (see Fig. 2), yet in this experiment the modulation of the adducin-AF647 labeling could affect the spectrin-CF680 pattern, but not the other way around. It might be due to a consistently higher localization density for AF647 staining compared to CF680, as we used secondary antibodies bearing 3-4 AF647 fluorophores, whereas the secondaries conjugated with CF680 had only a single conjugated fluorophore on average.

**Figure 4:**
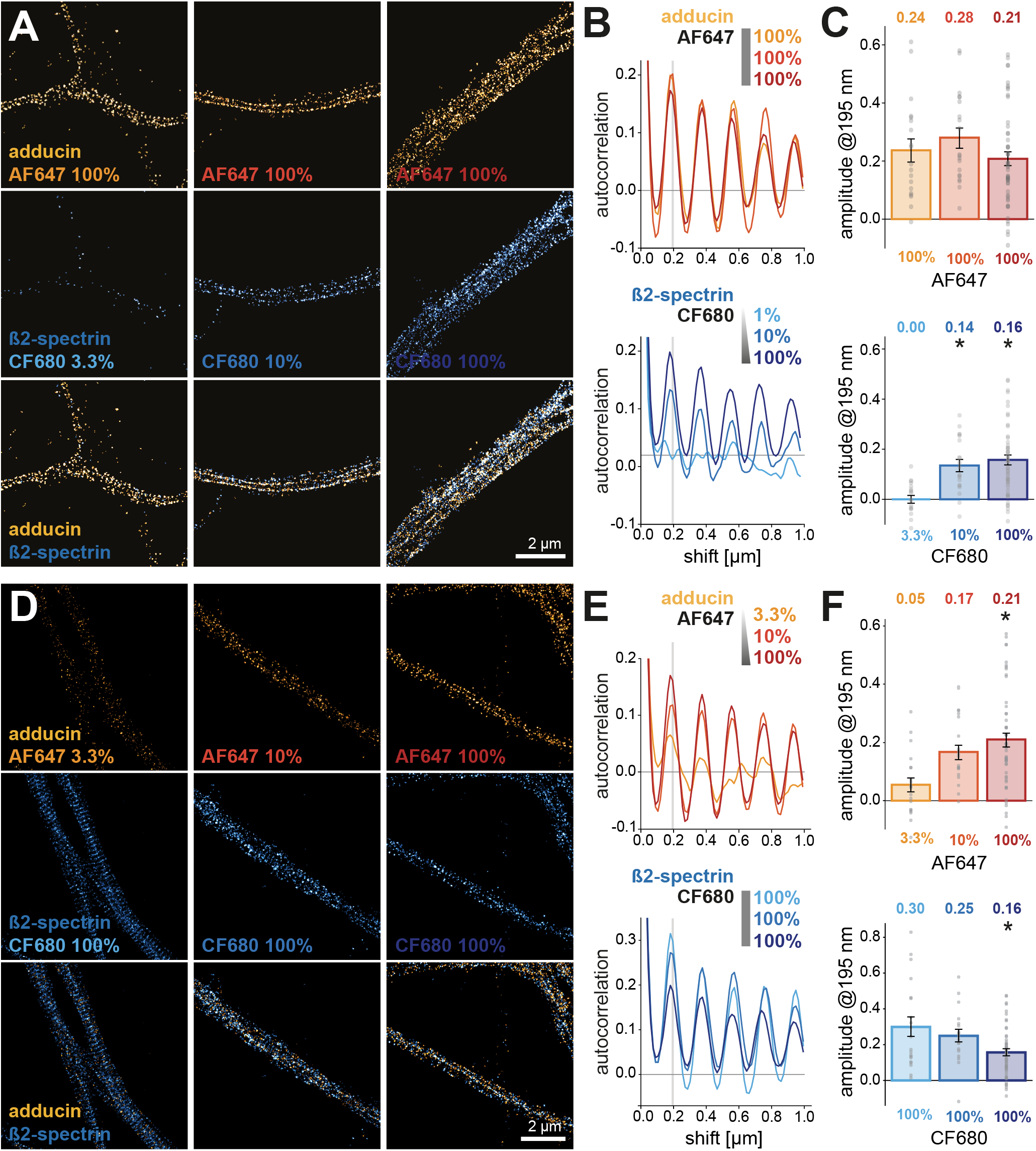
Effect of crosstalk on the detection of biologically-relevant patterns by SD-STORM. **A.** Reconstructed images from axons of hippocampal neurons stained for adducin (orange, top row) and ß2-spectrin (blue, middle row), imaged by SD-STORM using secondary antibodies conjugated with AF647 and CF680, respectively. During immunolabeling, the concentration of AF647-conjugated antibody was kept constant, while the concentration of CF680-conjugated antibody was increased successively from 3.3% to 10% and 100% (columns) of the total secondary antibody concentration (complemented with unlabeled secondary antibody). **B.** Autocorrelation curves from 1-μm long intensity profiles along axons for the constant adducin-AF647 (orange, top) and varying ß2-spectrin-CF680 (bottom, blue) channels for each secondary antibody concentration conditions. **C.** Amplitude of the autocorrelation first peak (base-subtracted height at 195 nm shift) for each secondary antibody concentration value. Dots are individual axonal segments, bars are SEM. Stars show a significant difference with the 100%-1% condition (first bar) by post-hoc Tukey-Kramer test, p<0.05. **D.** Converse experiment where the adducin imaging was done under varying AF647-conjugated secondary antibody concentration (from 3.3% to 100%, orange, top row) while ß2-spectrin imaging was done at constant 100% CF680-conjugated secondary antibody concentration (blue, middle row). **E.** Autocorrelation curves from 1-μm long intensity profiles along axons for the varying adducin-AF647 (orange, top) and constant ß2-spectrin-CF680 (bottom, blue) channels for each secondary antibody concentration conditions. **F.** Amplitude of the autocorrelation first peak (base-subtracted height at 195 nm shift) for each secondary antibody concentration value. Dots are individual axonal segments, bars are SEM. Stars show a significant difference with the 100%-1% condition (first bar) by post-hoc Tukey-Kramer test, p<0.05.

### Both S2C-PAINT and SD-STORM are readily compatible with astigmatism-based 3D SMLM

So far, we evaluated the crosstalk between channels for S2C-PAINT and SD-STORM, and its effect on biological pattern detection using 2D SMLM data. We next assessed if the two approaches can be combined with astigmatism PSF shaping, the most common approach to obtain 3D SMLM data allowing to add an axial coordinate to the blinking events localized within the imaged plane (Huang et al., 2008; Hajj et al., 2014; Diezmann et al., 2017). This is realized simply by inserting a cylindrical lens in front of each camera (Fig. S1) and performing Z calibration by moving fluorescent beads through the focal plane. We evaluated both S2C-PAINT and SD-STORM approaches using COS cells labeled for microtubules and clathrin (Jimenez et al., 2020). For S2C-PAINT, we used secondary antibodies conjugated to F1 docking strands to label microtubules, and to F2 docking strands to label clathrin, and imaged each target using IF1-Atto643 and IF2-Cy3B imagers (Fig. 5A). We obtained good structural quality for both channels, and Z color-coded single channel images (Fig. 5A, insets) show that we can get 3D localization over up to a ~1 μm range. Ratiometric analysis shows that simultaneous 2-color, 3D acquisitions yield similar results to the 2D case (see Fig. 1A) with most events localized only on one camera, allowing for a straightforward separation between channels (Fig. 5B). To visualize the 3D precision of our acquisition, we generated XZ sections along three microtubules and registered them to obtain an average microtubule profile (Fig. 5C, inset). We also calculated the microtubule section full width at half-maximum (FWHM) in the X and Z directions and found it to be 68 and 110 nm, respectively (Fig. 5C).

**Figure 5:**
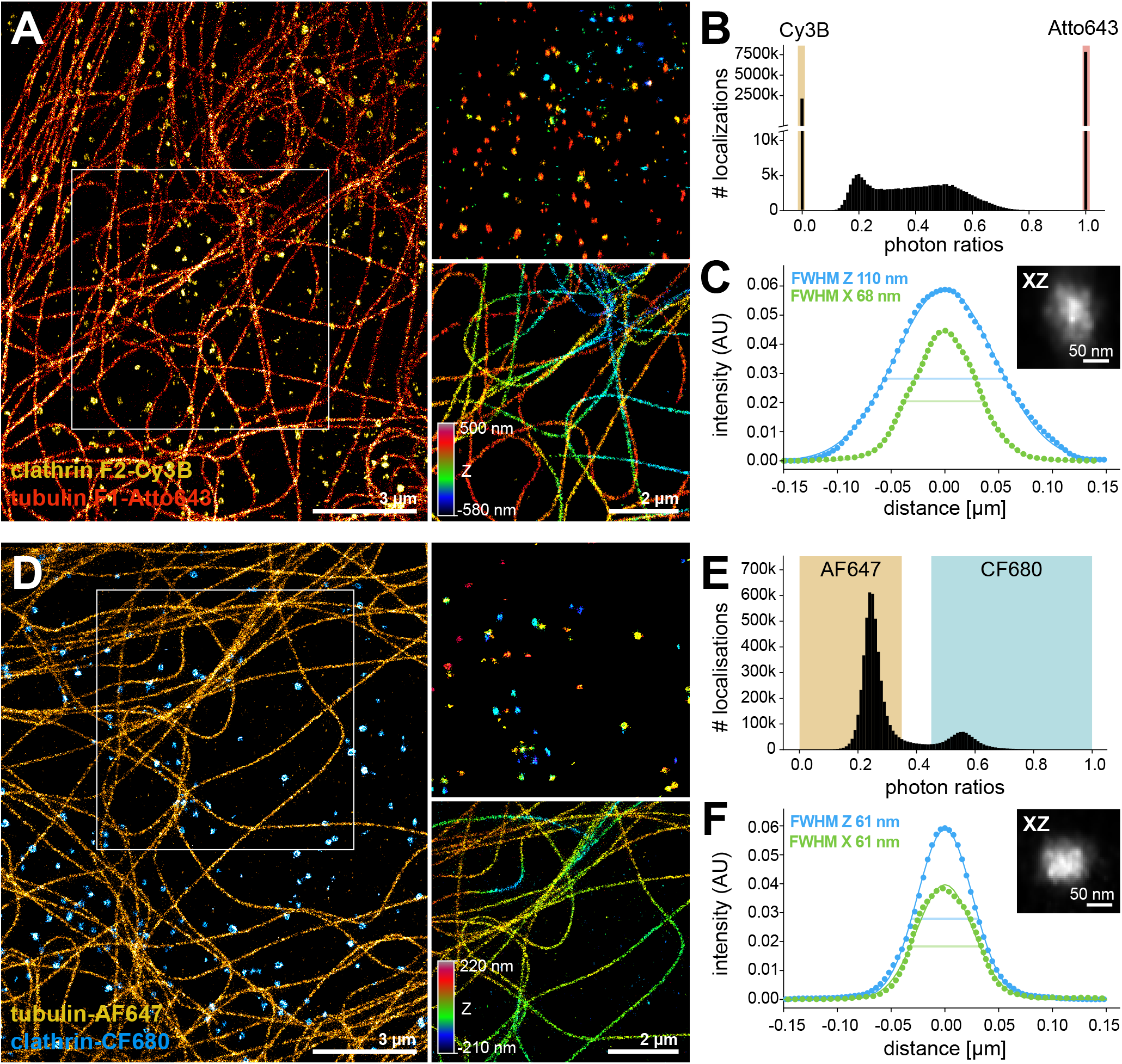
Extension of S2C-PAINT and SD-STORM to astigmatism-based 3D SMLM. **A.** 3D S2C-PAINT: image of a COS cell labeled for clathrin and tubulin, imaged with IF2-Cy3B (orange) and IF1-Atto643 (red), respectively. Insets show zoomed isolated channels, color-coded for Z. **B.** Distribution of the photon ratios for the acquisition in A, with colored areas for the ratios chosen for demixing: 0-0.01 for Cy3B (orange), 0.99-1 for Atto643 (red). **C.** Full-width half-maximum (FWHM) analysis of the intensity profile for transverse section along three isolated microtubules in A. The intensity profiles were fitted with a gaussian distribution for alignment and the width at half of the maximum in the X and Z direction was calculated. Inset, average transverse section obtained after alignment of individual sections. **D.** SD-STORM: image of a COS cell labeled for tubulin and clathrin, revealed with secondary antibodies conjugated to AF647 (yellow) and CF680 (cyan), respectively. Insets show zoomed isolated channels, color-coded for Z. **E.** Distribution of the photon ratios for the acquisition in A, with colored areas for the ratios chosen for demixing: 0.01-0.38 for AF647 (yellow area), 0.42-0.99 for CF680 (blue area). **F.** Full-width half-maximum (FWHM) analysis of the intensity profile for transverse section along three isolated micro-tubules in A. The intensity profiles were fitted with a gaussian distribution for alignment and the width at half of the maximum in the X and Z direction was calculated. Inset, average transverse section obtained after alignment of individual sections.

For SD-STORM, we labeled microtubules using an AF647-conjugated secondary antibody and clathrin using a CF680-conjugated one (Fig. 5D). Like for S2C-PAINT, we could obtain good structural quality for both channels together with Z localization across the focal plane (Fig. 5D, Z color-coded insets). The pairing analysis provided a distribution of photon ratios similar to the 2D case (see Fig. 2A), with an AF647 peak at ~0.25 and a CF680 peak at ~0.55, allowing to use the previously determined ratio ranges to separate the two channels (Fig. 5E). Averaging and analysis of microtubule cross-sections yielded a FWHM of 61 nm in both X and Z directions (Fig. 5F). This value is lower than the average thickness of microtubules obtained by PAINT, which is consistent with studies that directly compared the two methods (Früh et al., 2021).

### SD-STORM extension to simultaneous 3-target imaging

The first studies that developed SD-STORM already demonstrated that an extension to more than 2 colors should be straightforward (Bossi et al., 2008; Testa et al., 2010; Baddeley et al., 2011), with a progressive refinement toward the use of three far-red fluorophores (AF647, CF660C and CF680) excited by a 640-647 nm laser (Favuzzi et al., 2017; Andronov et al., 2022; Li et al., 2022). We thus extended our SD-STORM analysis to 3-color imaging using these fluorophores (Fig. 6), adding CF660C that has excitation and emission spectra with peaks in between those of AF647 and CF680 (Fig. 6A). As for the 2-color case, samples were illuminated with a single 640-nm laser and the emission of the fluorophore split by a dichroic mirror with a transition at 700 nm toward a reflected and transmitted pathway and their respective camera (Fig. S1).

**Figure 6:**
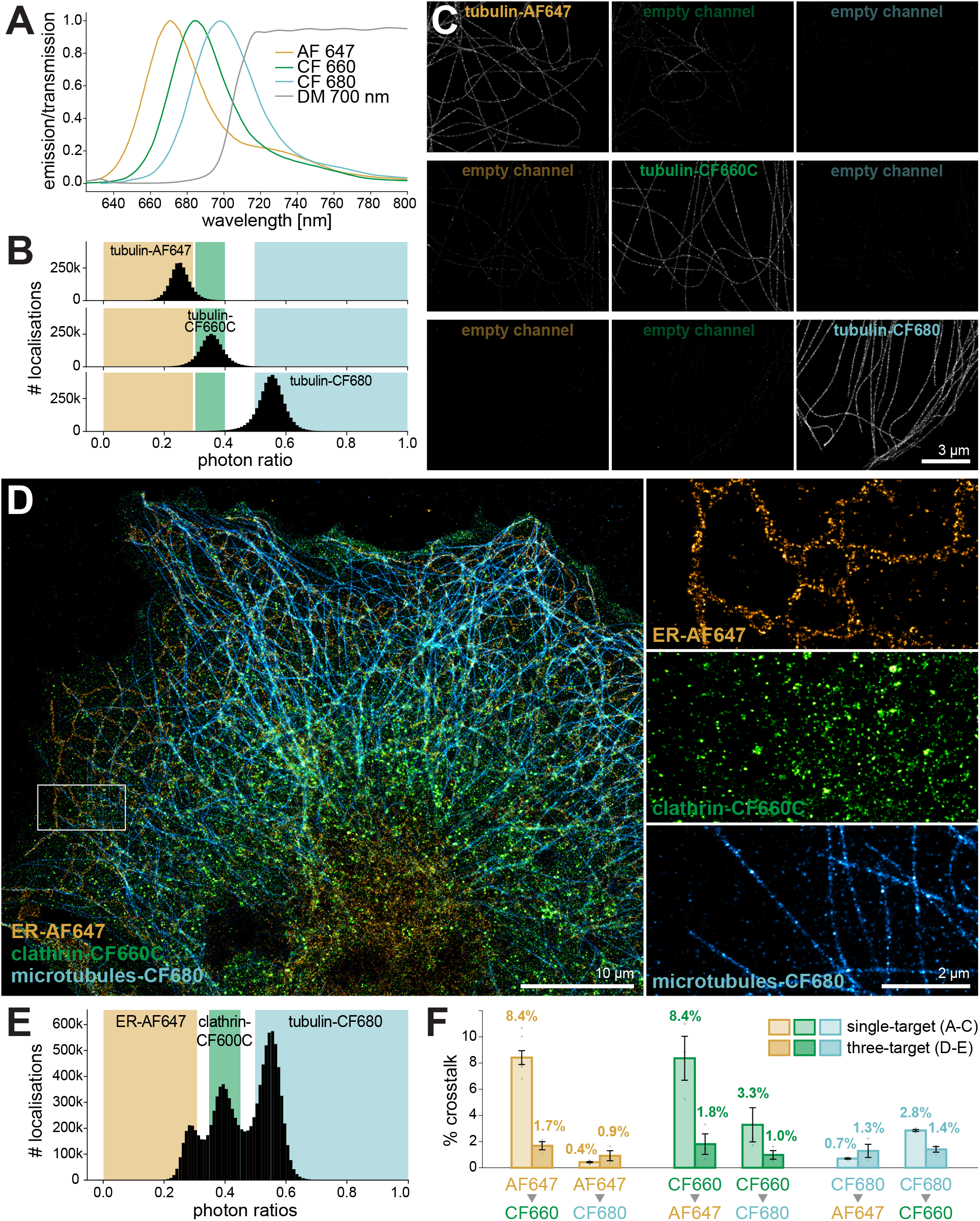
Extension of SD-STORM to 3 targets and crosstalk evaluation. **A.** Emission spectra of the fluorophores used for 3-color SD-STORM: AF647 (yellow), CF660C (green), and CF680 (blue), and transmission of the 700 nm long-pass dichroic inserted in the detection pathway (gray). **B.** Photon ratios for each fluorophore determined by ratiometric analysis from single-fluorophore stained microtubules in COS cells. Colored areas highlight the ratio ranges chosen for AF647 (0.01-0.29, yellow), CF660C (0.31-0.40, green), and CF680 (0.50-0.99, blue). **C.** SD-STORM images of COS cells stained for microtubules using secondary antibodies conjugated to AF647 (yellow, left column), CF660C (green, middle column), or CF680 (blue, right column). Reconstructions were performed in the three channels using the ratio ranges defined in B. ROIs segmented from the microtubule-channel were used to measure the number of localizations in each channel and calculate the crosstalk between channels (see F). **D.** 3-channel SD-STORM image of a COS cell labeled for the endoplasmic reticulum (ER, AF647), clathrin (CF660C), and tubulin (CF680). Insets show zoomed isolated channels. **E.** Photon ratios for each fluorophore from the 3-channel acquisition in D. Colored areas highlight the ratio ranges chosen to demix AF647 (0.01-0.30, yellow), CF660C (0.35-0.45, green), and CF680 (0.50-0.99, blue). **F.** Crosstalk between the channels calculated from the single-fluorophore staining (C) and the three-target staining (D) cellular samples. For the three-target sample, exclusive ROIs were used to delineate regions containing one target, but not the two others (see Fig. S5).

To assess the crosstalk between each channel, we used the two types of cellular samples previously validated for 2-color SD-STORM: COS cells labeled for microtubules using a single fluorophore (either AF647, CF660C or CF680, Fig. 5BC), and cells labeled for three targets using all fluorophores (Fig. 5D-E). Acquisitions from single-target labeled samples allowed us to obtain photon ratio distribution for each fluorophore, calculated as before as the ratio between the number of photons on the transmitted pathway image over the sum of photons detect on both images. AF647 and CF680 still had peaks at photon ratios of ~0.25 and 0.55, respectively, and CF660C peak was at a photon ratio of ~0.35 (Fig. 6B). From these distributions, we defined ratio ranges for each channel: 0.01-0.29 for AF647; 0.31-0.40 for CF660C, and 0.50-0.99 for CF680 (Fig. 6B). From the resulting single-color images (Fig. 6C), we could visually discern if some crosstalk was occurring, as well as measure the number of localizations in the non-target channels from ROIS defined on the target channel to calculate crosstalk values (Fig 6F). We found that AF647 crosstalk into CF660C and CF680 was 8.42% and 0.42%, respectively; that CF660C crosstalk into AF647 and CF680 was 8.36% and 3.28%, respectively; and that CF680 crosstalk into AF647 and CF660C was 0.70% and 2.85%, respectively (Fig. 6F). We then turned to simultaneous imaging of three targets using COS cells labeled for the endoplasmic reticulum (ER, anti-rtn4 antibody) with AF647, clathrin with CF660C, and microtubules with CF680 (Fig. 6D). As expected, the three photon ratio distribution were localized as in the single-target experiment, now overlapping significantly (Fig. 6E). For these 3-color acquisitions, we found that the photon ratio peak for CF660C could shift by up ~0.05 depending on the acquired sequence, and we accordingly adjusted the photon ratio ranges for demixing. Single-channel images demonstrate the good structural quality of the images for each structure (Fig. 6D, right). The same exclusive ROIs strategy as in the 2-color case, this time combining the three targets, was used to estimate the crosstalk directly from 3-color images. Interestingly, these led to values significantly lower than the single-target experiments: we found that AF647 crosstalk into CF660C and CF680 was 1.68% and 0.92%, respectively; that CF660C crosstalk into AF647 and CF680 was 1.80% and 0.98%, respectively; and that CF680 crosstalk into AF647 and CF660C was 1.29% and 1.40%, respectively (Fig. 6E). Despite these lower values, the crosstalk estimation obtained from 3-color images was consistent, with similar values in both directions for each fluorophore pairs, and considering the photon ratio distribution where the AF647 and CF660C are closer to one another (higher crosstalk) than CF660C and CF680 (lower crosstalk).

## Discussion

In this work, we implemented two approaches to perform simultaneous multicolor SMLM based on a module that splits emitted fluorescence toward two cameras. The first one consists of straightforward splitting the emission of two spectrally distinct red and far-red fluorophores, each excited by one laser. Interestingly, we found that this approach was better suited for simultaneous 2-color PAINT imaging (S2C-PAINT) than 2-color STORM: despite numerous attempts at finding good red fluorophores, such as Cy3B (Dempsey et al., 2011) or CF568 (Lehmann et al., 2016), their blinking characteristics are still not as good as far-red fluorophores, resulting in lower image quality (see Fig. S2). Multicolor PAINT has so far been performed almost exclusively by single-color sequential imaging known as Exchange-PAINT (Jungmann et al., 2014), but we reasoned that it should be readily amenable to simultaneous 2-color imaging using a mixture of imagers conjugated to red and far-red fluorophores, as we demonstrate here together with a couple of recent studies (Cheng et al., 2021; Chung et al., 2022). The second approach is based on demixing two far-red fluorophores excited by a single laser, with blinking events appearing with a different balance of intensities on each camera after splitting by a well-chosen dichroic mirror. This spectral demixing STORM (SD-STORM) method has been devised in the early days of SMLM (Schönle and Hell, 2007; Bossi et al., 2008; Testa et al., 2010; Baddeley et al., 2011; Gunewardene et al., 2011; Lampe et al., 2012), and it has since become a favorite option for home-made single-molecule microscope builders and users (Favuzzi et al., 2017; Gorur et al., 2017; Zhou et al., 2019; Sabinina et al., 2021) as it allows for 2-3 color imaging with good quality across channels and robustness to chromatic aberration and noise (Lampe et al., 2015). In our case, splitting the emission on two full-frame cameras allows for a large and flexible field-of-view, in combination with the illumination homogeneity ensured by an Adaptative Scanning for Tuneable Excitation Regions (ASTER) scanning module (Mau et al., 2021).

Despite its advantages, SMLM users often have the perception that SD-STORM would be more susceptible to crosstalk between channels than the classical 2-color approach. Therefore, we set up to rigorously measure the crosstalk of both methods using three different types of samples. The first type consists of two DNA origami nanorulers of different sizes seeded on the same coverslip (Lin et al., 2019; Scheckenbach et al., 2020). The idea was to create a sample where each target would be fully separated spatially and identifiable by its shape without relying on color/channel information. In theory, this would provide an unbiased assessment of crosstalk from a standard multicolor acquisition. Interestingly, the nanoruler samples provided higher values of crosstalk than cellular samples for both S2C-PAINT and SD-STORM (up to ~3.4% for the crosstalk of Atto655 into Cy3B in S2C-PAINT, see Fig. 1). This might stem from the low number of blinking events in the nanoruler acquisitions compared to the cellular staining experiments: the number of localizations within a nanoruler is much smaller than those within a single cellular structure, making the measurement more sensitive to noise and to spurious binding of the wrong imager strand. The nanoruler-based crosstalk values should thus be considered as a pessimistic estimate or higher boundary of the crosstalk, as in the case of a sample with a low density of labeling or higher background noise during the acquisition.

The second type of samples are cells labeled for a single target using each of the fluorophores used. This is a straightforward and widely-used approach to estimate crosstalk. However, when evaluating the S2C-PAINT approach where blinking event appear only on one camera (see Fig. 1), we realized that state-of-the-art SMLM algorithms based on an adaptive thresholding to detect blinking events introduce a massive bias in this approach, detecting a lot of background fluctuation events in the channel with no signal. This is particularly true for PAINT, where the presence of the diffusing imager strands generates a higher background. We thus devised a new crosstalk evaluation procedure for S2C-PAINT, taking advantage of the possibility to perform successive acquisitions of the same field of view with different imager mixtures. We were thus able to measure the change induced in a non-target channel by the presence of signal in the other channel, while compensating each for background. This approach yielded very low values for the S2C-PAINT crosstalk between Cy3B and Atto643 (0.4% and 0.02% for each direction, see Fig. 1). We interpret this result as the most exact for S2C-PAINT, as we were able to use true multicolor acquisitions and to compensate for the background and adaptive nature of the blinking event detection algorithm. For SD-STORM, blinking events appear on both cameras, allowing for a straightforward single-target strategy even when using an adaptive algorithm. As for S2C-PAINT, the single-target samples showed the lowest crosstalk across samples for AF647 and CF680 (0.7% and 0.5% for each direction). Finally, we attempted to evaluate the crosstalk on 2-target cellular samples, to determine if a reliable estimate could be directly obtained from multicolor samples. Using well-segregated cellular structures (microtubules and clathrin-coated pits) and a subtractive strategy to generate “microtubule-only” and “clathrin-only” ROIs, we obtained crosstalk estimations that are consistent with the two other samples. Values were higher that the single-target sample estimation, which might be caused by residual overlap between structures, or the non-perfect specificity of the primary and secondary antibodies. Single-color sample estimation might be more accurate, but a direct measurement of crosstalk on a multicolor sample can provide meaningful higher boundaries of the crosstalk values.

Overall, the three types of samples provide different crosstalk values, likely due to subtle differences in what exactly is measured. This stresses the importance of detailing the type of sample and analysis used to determine crosstalk values across hardware configurations or microscope setups. In particular, the estimation using single-target samples should be performed carefully when using adaptive SMLM algorithms. Despite those difference in absolute values, we found consistent trends in the crosstalk pattern across samples and methods. For S2C-PAINT and 2-color SD-STORM, we consistently found the crosstalk from the higher wavelength channel into the lower wavelength channel to be higher than in the reverse direction. As the ratiometric analysis should help removing blinking events with unbalanced photon ratio (Lampe et al., 2015), it is likely that the crosstalk localizations are related to non-specific elements in the staining, imaging or processing rather than true optical bleed-through from the theoretical fluorophore spectra and filter curves: non-specific binding of imager strands in S2C-PAINT, pairing errors in SD-STORM. It is likely that our efforts in minimizing crosstalk and bias that result in quite low values compared to the literature stem from a combination of residual factors that will be difficult to eliminate further. Refinements in channel assignment in SD-STORM (Andronov et al., 2022; Li et al., 2022; Siemons et al., 2022), or more complex techniques such as spectral STORM (Yan et al., 2018) might allow for an even better accuracy in this regard. It must be noted that the crosstalk values we obtained from 2-color SD-STORM are only slightly higher than those obtained for optimized S2C-PAINT, demonstrating the robustness of SD-STORM to crosstalk.

In addition to evaluating the crosstalk on different types of samples, we also tried to evaluate how it could affect the results of experiments by interfering with a known biological pattern. Usually, crosstalk is evaluated by comparing the same target across channels (single-target samples) or target with similar abundance (microtubule and clathrin in our 2-target samples). However, crosstalk is likely to be more detrimental in experiments where one target has a low abundance compared to the other. To assess this, we took advantage of neuronal axons where two proteins (ß2-spectrin and adducin) show a similar labeling density with a complementary, 190-nm periodic staining pattern (Xu et al., 2013; Leterrier, 2021). We could assess if modulating the labeling density of one target would affect the other target. Interestingly, we did not detect an impact of a varying labeling density in one channel on the other channel in S2C-PAINT, but could see a slight drop in the periodicity for CF680-labeled ß2-spectrin when rising the labeling density of AF647-labeled adducin (see Fig. 4). As CF680 blinking events are usually less numerous (due to the lower degree of labeling of the CF680-conjugated secondary antibodies) and less bright than AF647 blinking events, it is possible that this condition allowed us to unmask how crosstalk can impact the detection of a sub-diffraction pattern in a biologically-relevant experiment.

Finally, we confirmed that both S2C-PAINT and SD-STORM can readily be extended to astigmatism-based 3D SMLM, with no alteration in the image quality or channel separation (see Fig. 5), and showed how SD-STORM can be extended to 3 colors by using CF660C, a fluorophore that sits in-between AF647 and CF680 (Favuzzi et al., 2017; Andronov et al., 2022; Li et al., 2022). This results in higher crosstalk between the channels, as the overlap in photon ratio becomes significant (see Fig. 6). However, one advantage of SD-STORM is that the specificity can be optimized by tuning the photon ratio ranges for each channel, to the cost of more rejected localizations and a sparser image. One could imagine extending S2C-PAINT to three colors, introducing a third fluorophore-conjugated imager strand with an emission spectrum that would be split by the 662 nm dichroic toward each camera (such as Atto594), rather than being sent to only one, adding a component of spectral demixing to the 2-color separation approach. Spectral demixing with PAINT has recently been demonstrated with three far-red fluorophores, to the cost of lower maximal blinking density, as blinking events from all three fluorophores appear on both cameras (Gimber et al., 2022).

More generally, a prevalent disadvantage of SMLM is its low throughput: long acquisition times drastically limit the number of experiments that can be performed, as well as making the imaging process more susceptible to drift. Optimizing the acquisition speed is thus an important progress area for STORM and particularly for PAINT which is even slower (Schueder et al., 2019; Civitci et al., 2020; Strauss and Jungmann, 2020). One way to optimize the “localization throughput” of an SMLM experiment is to get as close as possible to the maximum blinking density that the processing algorithm can handle, with higher fractions of active fluorophores in STORM or higher binding rates of imager strands in PAINT. In this regard, spectral demixing strategies are dividing the maximal attainable density by the number of fluorophores imaged, as they all appear on each camera frame. Yet for the two optimized strategies we tested here, 2-color SD-STORM is not slower than S2C-PAINT, because the loss in maximum density is more than compensated by the faster blinking and lower exposure times possible with STORM compared to PAINT. For 2-color acquisitions, we thus would favor SD-STORM over S2C-PAINT as it is compatible with initial observation before super-resolved acquisition, faster, straightforward to implement with a commercially available setup (Jackson et al., 2022; Tomer et al., 2022; Gazzola et al., 2023) and readily extensible to 3-color imaging. A key advantage of S2C-PAINT in the future is that implementing successive rounds of PAINT imaging would provide a straightforward way to extend it to multiplexing with 4 or more channels, with a significant gain in speed compared to single-color Exchange PAINT.

## Acknowledgments

We ackowledge funding by the *Agence Nationale pour la Recherche* (ANR-20-CE16-021-03 to C.L. and ANR-21-CE42-0015 to N.B., S.LF and C.L.), and the *Association Nationale de la Recherche et de la Technologie* (ANRT) for the co-funding of K.F. (CIFRE PhD). This work has received funding from the European Union’s Horizon 2020 research and innovation programme under grant agreement #871124 Laserlab-Europe (JRA ALTIS). We thank Fanny Boroni-Rueda for neuronal cell cultures, and Marie-Jeanne Papandréou for discussions.

## Author contributions

Conceptualization: K.F., N.B, S.L.F, C.L.; Methodology: all authors; Formal analysis: K.F.; A.M.; Investigation: K.F.; Resources: V.C., N.B.; Writing-Original Draft: K.F., C.L.; Writing-Review and Editing: all authors; Visualization: K.F., C.L.; Supervision, Project Administration: C.L, Funding acquisition: N.B., S.L.F., C.L.

## Methods

### COS cells cultures and fixation procedure

COS-7 cells (ATCC CRL-1651) were cultured in DMEM medium (ThermoFisher #61965026), supplemented with fetal bovine serum (ThermoFisher, #A3381901) and antibiotics (Penicillin/Streptomycin, ThermoFisher #15140122). 24 hours after seeding on poly-L-lysine (Sigma #P2636) coated coverslips (#1.5H 18 mm Marienfeld, VWR) to a density of about 10%, they were either extracted and fixed with a two-step protocol using glutaraldehyde as a fixative (Sigma #3G5882), or fixed in a single step with a mixture of glutaraldehyde and paraformaldehyde (PFA, Delta Microscopy #EM-15714) For the two-step extraction/fixation with glutaraldehyde, cells were first extracted by a 45-second incubation with 37°C pre-heated 0.1% glutaraldehyde, 0.25% triton (Sigma #T8787) in PEM buffer (80mM PIPES, 2mM MgCl2, 5mM EGTA, pH 6.8) then fixed for 10 minutes at 37°C in pre-heated 2% glutaraldehyde, 0.5% Triton in PEM. For the single step glutaraldehyde/PFA fixation, cells were fixed for 10 minutes at 37°C with 0.1% glutaraldehyde, 4%PFA, and 4% (w/v) sucrose in PEM buffer. Following both fixation procedures, cells were rinsed in phosphate buffer (PB, 0.1 M pH 7.3), then residual glutaraldehyde was quenched for 7 minutes using 10 mg/ml sodium borohydride (Sigma #213462) in PB before further rinses with PB (Jimenez et al., 2020).

### Neuronal cultures

All procedures involving animal cell culture followed the guidelines from European Animal Care and Use Committee (86/609/CEE) and were approved by the Aix-Marseille university ethics committee (agreement D13-055-8). Rat hippocampal neurons were cultured on top of a glia feeder layer according to the Banker protocol (Kaech and Banker, 2006). Briefly, hippocampi from Wistar rat embryos (Janvier Labs) were dissected, then cells were homogenized and seeded on poly-L-lysine coated #1.5H glass coverslips to a density of 4000 cells per cm2 in MEM (ThermoFisher #21090-055) supplemented with fetal bovine serum, which was replaced after three hours with Neurobasal medium (ThermoFisher #21103-049) supplemented with B27 (Thermo Fisher #17504-044). Mature neurons were fixed after 14 days in culture using 4% PFA and 4% sucrose in PEM buffer for 20 minutes at room temperature (RT) and rinsed with PB.

### Immunostaining and antibodies

Blocking and permeabilization were performed in immuno-cytochemistry buffer (ICC: 0.22% gelatin, 0.1% Triton X-100 in PB) for 1 to 3 hours on a rocking table. Cells were incubated overnight at 4 °C with primary antibodies diluted in ICC, rinsed, and incubated with secondary antibodies diluted in ICC for 1 hour at RT. After final rinses in ICC and PB, the samples were stored in PB with 0.02 % (v/v) sodium azide (Sigma #08591) before imaging. Primary and secondary antibodies used are summarized in Table S2 and S3.

### SMLM microscope

The SMLM microscope used for all imaging is based on either an Olympus IX81 inverted microscope stand equipped with a focus stabilization system (ZDC2, Olympus) and a 100X, NA 1.49 oil objective (APON100XHOTIRF, Olympus) or a Nikon Ti2 inverted microscope stand equipped with a motorized stage, a piezo Z-stage (Mad City Labs), a focus stabilization system (Perfect Focus System, Nikon) and a 100X, NA 1.49 oil objective (CFI SR HP Apochromat TIRF 100XC, Nikon). An Abbelight SAFe360 module is attached to the side C-mount of the stand. It receives laser excitation from an Oxxius L4Cc Combiner equipped with 640 nm (500 mW), 532 nm (400 mW), 488 nm (150 mW) and 405 nm (100 mW) lasers through the ASTER scanning module (Abbelight)(Mau et al., 2021). Separation between excitation and emission bands is done using a quad-band dichroic mirror (Semrock, Di03-R405/488/532/635-t3-25×36), filtered on both paths by quadband emission filters (Semrock FF01-446/510/581/703-25) and captured by two Hamamatsu Photonics Flash4 V3 sCMOS cameras. For 3D imaging, the point spread functions were shaped using two cylindrical lenses inserted in the optical pathway between the second dichroic mirror and the cameras.

### DNA-PAINT acquisition

For DNA-PAINT imaging, samples were mounted in an open metal chamber (Ludin Chamber, Life Imaging Services) allowing for easy medium exchange. The “regular” imager strands (I1 and I3)(Jungmann et al., 2014) or repetitive-sequence “fastPaint” imager strands (IF1 and IF2) (Strauss and Jungmann, 2020) were diluted in imaging buffer and washed off with washing buffer (imager strands and buffers from Massive Photonics). Fluorescence illumination was performed using the 532 nm and 640 nm lasers in Highly Inclined and Laminated Optical sheet (HILO) illumination to restrict illumination to ~1 μm above the coverslip for minimal background fluorescence from unbound imagers. The illumination strength was adapted to the field of view we chose, between 50 μm × 50 μm and 80 μm × 80 μm. With using 40-100 % of the laser, the irradiance resulted in 0.55-3.44 kW/cm2. The emission was split in a reflected and a transmitted pathway by a 662 nm dichroic mirror (Semrock, FF662-FDi02-t3-25×36); acquisitions were acquired with exposure times between 30 and 100 ms. A detailed table of the parameters for each acquisition is provided in Table S1.

### Spectral demixing-STORM acquisition

STORM samples were mounted in sealed silicone chambers filled with Abbelight STORM buffer kit. The samples were labeled with secondary antibodies conjugated with AF647 and CF680 (see table 2) for two-target spectral demixing; for three targets a staining with an antibody conjugated with CF660C was added (see Table above). Illumination was performed using the 640 nm laser (Between 1.23 and 13.62 kW/ cm2 depending on the field of view) with manual increase of low-power illumination using the 405 nm laser line aid fluorophore recovery from long-lived dark states. HiLO illumination was used to restrict illumination to ~1 μm above the coverslip. The detection pathway was split into a reflected and a transmitted part by a 700 nm dichroic mirror (Semrock, FF699-Fdi01-t3-25×36). Exposure time was chosen depending on the size of the field of view between 5 ms (30 um μm × 30 μm field of view) and 50 ms (100 μm × 100 μm field of view); and between 15,000 and 60,000 frames were acquired. All parameters per acquisition are summarized in Table S1.

### Nanorulers sample and imaging

The DNA origami-based nanorulers slide was custom ordered (Gattaquant) and contains two types of 3-spot nanorulers deposited in a single fluidic cavity allowing for medium exchange and rinses. One nanoruler type has spots spaced by 40 nm tagged with a P1 docking strand, the other one has spots spaced by 80 nm tagged with a P3 docking strand. For simultaneous 2-color DNA-PAINT, imager strands conjugated with Cy3B and Atto655 (all imagers from Massive Photonics) were diluted in phosphate buffer saline (PBS) + 10 mM MgCl2 to a concentration between 500 pM and 20 nM. The sample was simultaneously illuminated in HiLO with 532 nm (0.37-0.71 kW/cm2) and 640 nm (0.91-1.77 kW/cm2) lasers, emission was split using the 662 nm dichroic mirror and recorded 60k frames with an exposure time of 50 ms on the two cameras on a 70 μm × 70 μm field of view.

For far-red spectral demixing DNA-PAINT, 3 to 10 nM I1-Atto680 and 2 nM I3-Atto655 were used to reveal the nanorulers in PBS + 10 mM MgCl2; the sample was illuminated with a single 640 nm laser (2.5 kW/cm2 in HiLO and emission was split with the 700 nm dichroic mirror. The cameras were set to an exposure time of 100 ms and to record 60,000 frames.

### SMLM data processing

Acquired PAINT and STORM sequences were processed using the Abbelight Neo Analysis software. Detection of intensity peaks used a wavelet algorithm (Izeddin et al., 2012) after local means background estimation and removal. Intensity peaks at least twice as high as the background with a size of 3×3 to 7×7 pixels area (300 to 700 nm) were considered a single-molecule blinking event and further processed for fitting using gaussian fitting with least-squares error. The number of photons emitted by the blinking event was estimated from the background-subtracted raw data by integration over a 11×11 pixel round area (1.1 μm diameter)(Bourg et al., 2015). Frame sequences from each camera were processed individually to generate two lists of localizations containing their coordinates and photon number. Three-dimensional astigmatism-based acquisition were fitted for Z position according to the eccentricity of the PSF using a calibration obtained on 100 μm beads.

### Channel assignment (demixing) from two-camera data

After fitting, two images were reconstructed using the localizations list from each camera and were then aligned using an affine transform. The localization coordinates obtained from each camera image sequence were then modified using the determined affine transformation, before checking for pairs of localizations between the two cameras: localizations appearing at the same coordinates with a tolerance of 500nm on identical frames. For each detected pair, the ratio of the emitted photons was calculated with r=I_trans_/(I_trans_+I_ref_) where Itrans is the number of photons for the localization on the transmitted path camera image, and I_ref_ the number of photons for the localization on the reflected path camera image. Unpaired localizations only appearing on the reflected or transmitted path camera image were assigned a ratio of 0 or 1, respectively. For S2C-PAINT data, channels were either directly defined from the localization files from each camera, or after demixing. In the latter case, localizations with ratios 0-0.01 were assigned to the Cy3B/Atto565 channel, and localizations with rations 0.99-1 were assigned to the Atto643 channel. For 2-color SD-STORM data, localizations were always demixed: those with ratios 0-0.29 were assigned to the AF647 channel, and those with ratios 0.5-1 were assigned to the CF680 channel. For 3-color SD-STORM data, localizations with ratios 0.01-0.29 were assigned to the AF647 channel, 0.31-0.45 to the CF660C channel, 0.5-0.99 to the CF680 channel. The boundaries varied slightly between experiments, for repetitions of the same experiment, the exact same boundaries were chosen. After demixing, separate localization files were generated for each channel, with the final coordinate of each localization determined using weighted averages of the localization precision of the localizations on the two cameras and these localization files were used to reconstruct images for each channel.

### Crosstalk analysis

#### Analysis of crosstalk from nanorulers images

Nanoruler analysis was performed on localization tables, namely the x,y positions, and the number of photons for each detection channels. To detect groups of localization Density Based Clustering (DBSCAN)(Ester et al., 1996) algorithms were used: from a starting localization point, they will iteratively include localizations closer than a typical distance ε, until no more neighboring candidates is found. First a DBSCAN was used to detect and filter out localization points with less than 4 neighbors on ∊=15 nm distances. These points are attributed to background or detection of noise and could not be associated to nanorulers. A second DBSCAN was used to isolate clusters of >7 localizations, on typical distances of ∊=80 nm. This results in isolated clusters for nanorulers of 40 nm and 80 nm spacing. For each nanoruler, we then adjusted the distribution of three spots by a Gaussian Mixture Model (GMM), which fits the localization cloud as arising from three normal distributions. GMM estimates the mean and standard deviation of each spot, allowing for size estimation and estimation of the localization precision. DBSCAN and GMM algorithms were implemented via the library scikit-learn (Virtanen et al., 2020). Crosstalk then calculated from the number of localizations in each channel (Fig. S3).

#### Analysis of multicolor DNA-PAINT crosstalk from single-target samples

DNA-PAINT crosstalk was evaluated using a twostep strategy with varying imagers (Fig. S4). A sample labeled for a single target and docking strand (example: microtubules with an F2-coupled secondary antibody) was first imaged with two non-target imagers (A1Ch1_NT: IF1-Atto565 and A1Ch2_NT: IF1-Atto643). After rinses, the same field of view was then imaged using a target and a non-target imager (A2Ch1_T: IF2-Atto565 and A2Ch2_NT: IF1-Atto643). Both acquisition sequences (A1 and A2) were processed using localization and optional demixing. Images for all channels from both acquisition sequences were reconstructed using the histogram method in ThunderSTORM (Ovesny et al., 2014), where each pixel (15 nm in size) takes a value corresponding to the number of localizations inside this pixel. A ROI for the target (microtubules) was obtained by thresholding the reconstructed image of the target imager from the second acquisition sequence (A2Ch1_T: IF2-Atto565) and used to measure the number of localizations in this channel (A2Ch2_T), then in all other channels from both acquisition sequences (A1Ch1_ NT, A1Ch2_NT and A2Ch2_NT). The crosstalk of the target channel (Ch1: Atto565) into the non-target channel (Ch2: Atto643) is then defined as the number of localizations in the non-target channel (Ch2: Atto643) that are added by the presence of a target imager in the target channel (Ch1: Atto565), corrected for the background obtained using non-target imagers in each channel (A1Ch2_NT and A1Ch1_NT), expressed in percentage∷

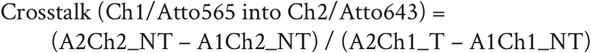

#### Analysis of demixing-STORM crosstalk from single-target samples

Sample were labeled for a single target (microtubules) with a given fluorophore, then imaged and processed according to the demixing-STORM procedure above. Localizations were then assigned to two (AF647 and CF680) or three (AF647, CF660C, CF680) channels using the demixing procedure detailed above, and the images from each channel was reconstructed using the histogram method. A ROI was obtained by thresholding the microtubules on the channel corresponding to the fluorophore used (target channel), and the number of localizations was measured inside this ROI on the images of the target channel and those of the non-target channel(s) using a dedicated Fiji macro (“Split Locs by ROI” macro from the ChriSTORM GitHub repository at https://github.com/cleterrier/ChriSTORM/). The proportion of localizations within the ROI in each channel - which is the crosstalk value for non-target channels - was obtained by dividing the number of localizations in this channel by the total number of localizations in all channels and was expressed as a percentage.

#### Analysis of multicolor DNA-PAINT and demixing-STORM crosstalk from multicolor samples

A similar procedure was used to estimate crosstalk on samples labeled for multiple targets. After imaging, processing and demixing into channels, ROIs were obtained for each channel by thresholding the corresponding reconstructed images. To avoid bias from target overlap (such as clathrin-coated pit overlapping with a microtubule), overlapping areas of ROIs from the different channels were excluded before counting the localizations within each ROI on the target and non-target images. The localization proportions (crosstalk value for non-target images) were then calculated as above.

### Analysis of the actin/spectrin periodicity along axons

The investigation of the effect of crosstalk at different labeling densities was performed on neurons labeled for ß2-spectrin and adducin, which form a complementary periodic scaffold with a 190 nm periodicity. After S2C-PAINT or SD-STORM imaging of the samples, the acquisition sequences were processed and demixed into two channels as described above. Linear segments of axons were manually selected and analyzed by autocorrelation for each channel using a custom script (“Autocorrelation” script available at the Process_Profile GitHub repository https://github.com/cleterrier/Process_Profiles), to determine if the presence of the periodicity observed in one channel could be perturbed by crosstalk from the presence of a second channel. This was done by subtracting the amplitude of the autocorrelation curve at the first valley (95 nm shift for a 15-nm pixel size) to its amplitude at the first peak (195 nm shift).

### Microtubule cross-section analysis

Microtubules from 3D acquisitions with S2C-PAINT and SD-STORM were analyzed for Full Width at Half Maximum (FWHM) in the X and Z dimension. Three microtubule sections were taken from each image and turned into perpendicular XZ cross-sections reconstructed with 4-nm pixel size (“Line ROIs to Slices” and “Generate Zooms and Slices” macros form the ChriSTORM GitHub repository). The intensity profile was taken from the whole width of the reconstructions and aligned (“ProFeatFit” script scrip available from the Process_Profile GitHub repository). The profiles were then averaged and fitted with a gaussian distribution and the FWHM calculated using scipy optimize functions.

## Supplementary Figures and Tables

**Figure S1:**
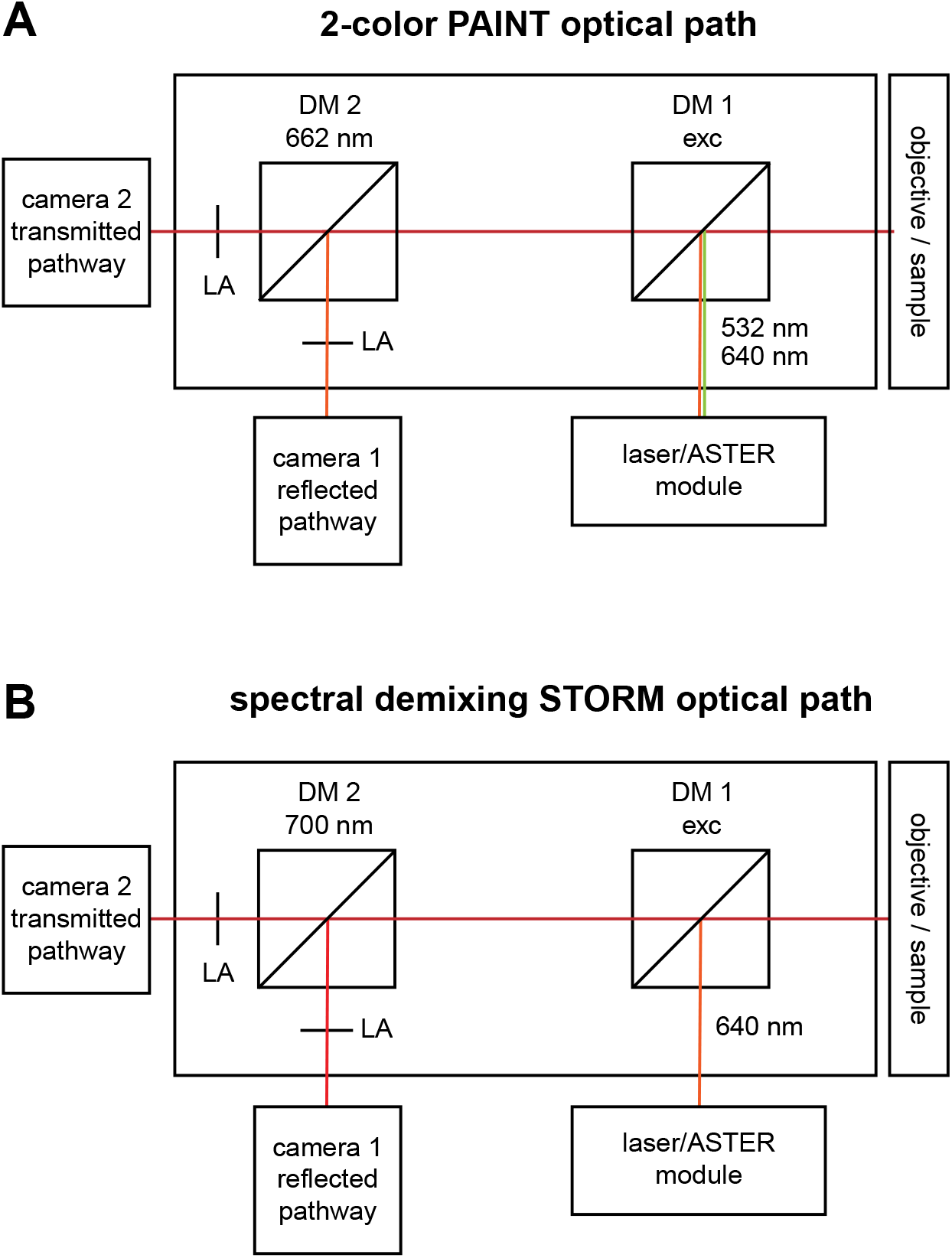
Optical paths for S2C-PAINT and SD-STORM. **A.** Optical path for 2C-PAINT: the 532 and 640 nm excitation laser beams are reflected by the dichroic mirror DM1 into the microscope body (objective and sample). The fluorescence emitted from the sample travels through DM1, and is split by the 662 nm dichroic mirror DM2 toward the two cameras. For 3D acquisitions, cylindrical lenses (LA) are inserted in front of each camera. **B.** Optical path for SD-STORM: the single 640 nm excitation laser beams is reflected by the dichroic mirror DM1 into the microscope body (objective and sample). The fluorescence emitted from the sample travels through DM1, and is split by the 700 nm dichroic mirror DM2 toward the two cameras. For 3D acquisitions, cylindrical lenses (LA) are inserted in front of each camera.

**Figure S2:**
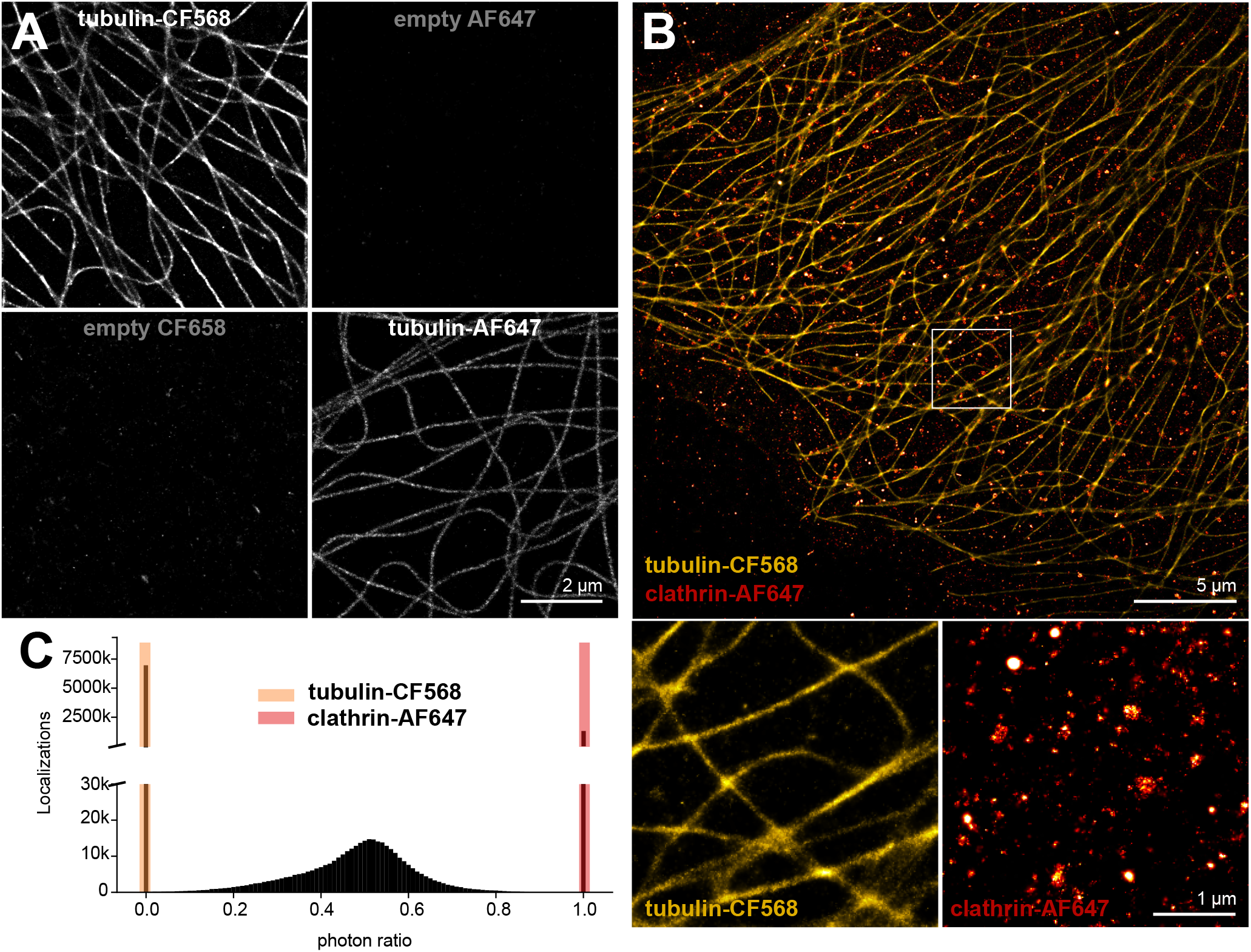
Simultaneous 2-color STORM with CF568 and AF647. **A.** COS cells stained for tubulin with a secondary antibody conjugated to CF568 (top row) or AF647 (bottom row), demixed into 2 channels with the 2-color ratio ranges (0-0.01 for CF568, 0.99-1 for AF647). **B.** Simultaneous 2-color STORM image of a COS cells stained for tubulin with a secondary antibody conjugated to CF568 (orange) and clathrin with a secondary antibody conjugated to AF647 (red), demixed into 2 channels with the 2-color ratio range (0-0.01 for CF568, 0.99-1 for AF647) defined in C. Bottom images are zoomed isolated channels. One can see the low quality of the microtubule image, due to the sub-optimal blinking properties of CF568. **C.** Ratiometric analysis for the image shown in B, with used 2-color ratio ranges highlighted in colors.

**Figure S3:**
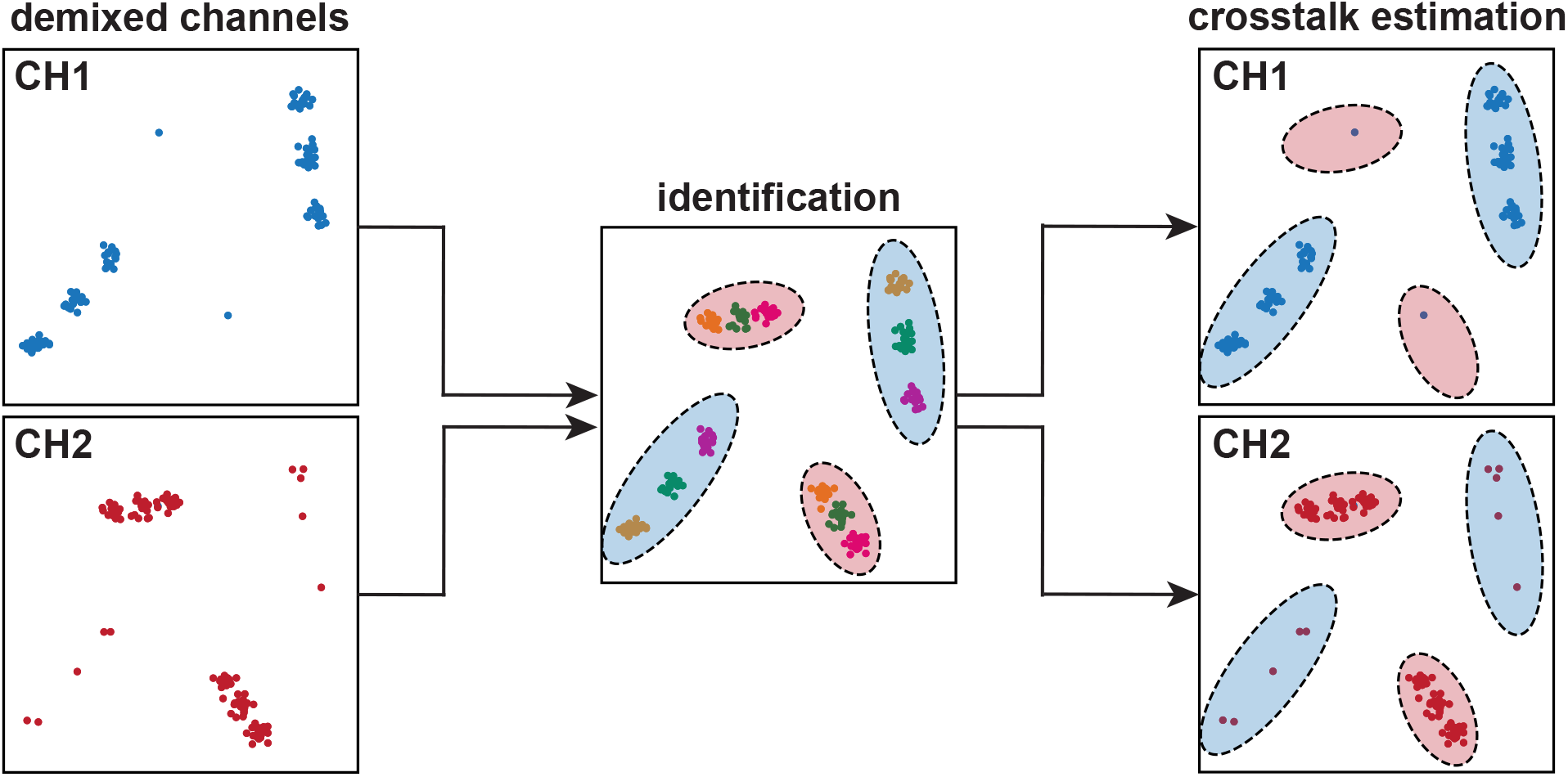
Crosstalk analysis for the nanoruler experiments. Two channels are obtained after acquisition and demixing with pre-determined ratio ranges (left column). The image is then processed with a small-distance DBSCAN (center panel) to segment the individual docking sites (colored clusters) and a large-distance DBSCAN to segment individual nanorulers (dashed ellipses). A Gaussian Mixture Model is used to fit the position of the docking sites and the total size of the nanorulers, resulting in a channel-independent classification in long and short nanorulers (blue and red ellipses in the right column). Finally, number of localizations inside each classified nanoruler is counted in each channel to calculate the crosstalk for each nanoruler (right column).

**Figure S4:**
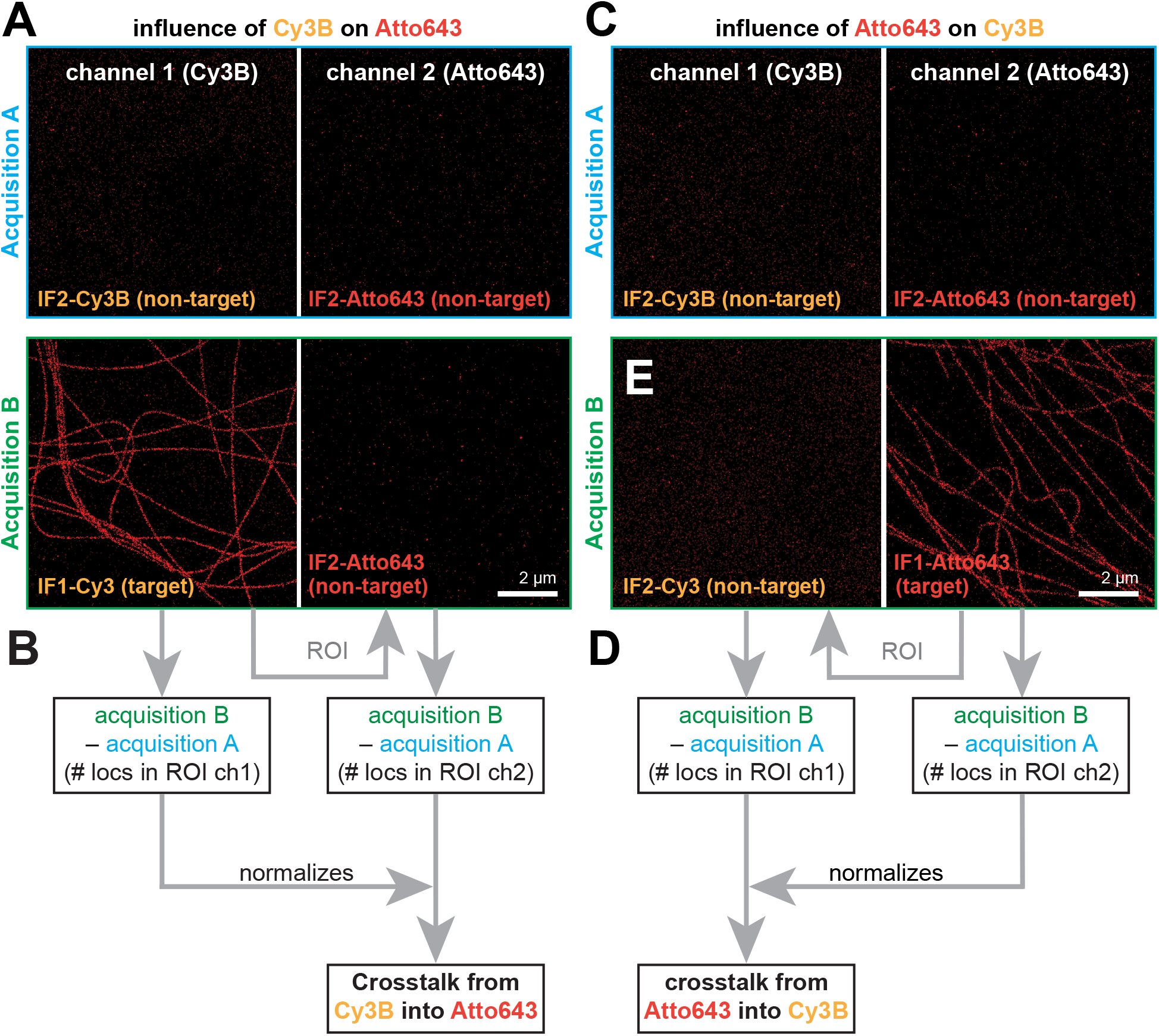
Crosstalk evaluation from single-target staining in S2C-PAINT based on successive acquisition of background and target images. **A-B.** Evaluation of the crosstalk from Cy3B (orange) into Atto643 (red), i.e., influence of a signal in the Cy3B channel on the Atto643 signal. **A.** COS cell stained for tubulin with a secondary antibody conjugated to an F1 docking strand is subjected to 2 successive acquisitions: in acquisition A, both channels contain a non-target imager (IF2-Cy3B and IF2-Atto643), resulting in background in both channels (first row, blue). In acquisition B, the Cy3B channel contains a target imager (IF1-Cy3B) resulting in signal, while the Atto643 channel contains a non-target imager (IF2-Atto643), resulting in background (green, bottom row). B. A ROI is drawn from segmenting the microtubule signal in the I1-Cy3B channel of acquisition B, and used to measure the number of localizations in all channels. Acquisition A is used to subtract background in both channels from acquisition B, and the remaining localizations in the Atto643 channel in acquisition B (assumed to be due to the presence of Cy3B signal) are normalized by the number of localizations in the Cy3B signal channel of acquisition B, resulting in the crosstalk value for Cy3B into Atto643. **C-D.** Evaluation of the crosstalk from Atto643 (red) into Cy3B (orange), i.e., influence of a signal in the Atto643 channel on the Cy3B signal. C. COS cell stained for tubulin with a secondary antibody conjugated to an F1 docking strand is subjected to 2 successive acquisitions: in acquisition A, both channels contain a non-target imager (IF2-Cy3B and IF2-Atto643), resulting in background in both channels (first row, blue). In acquisition B, the Cy3B channel contains a non-target imager (IF2-Cy3B) resulting in background, while the Atto643 channel contains a target imager (IF1-Atto643), resulting in signal (green, bottom row). **D.** A ROI is drawn from segmenting the microtubule signal in the I1-Atto643 channel of acquisition B, and used to measure the number of localizations in all channels. Acquisition A is used to subtract background in both channels from acquisition B, and the remaining localizations in the Cy3B channel in acquisition B (assumed to be due to the presence of Atto643 signal) are normalized by the number of localizations in the Atto643 signal channel of acquisition B, resulting in the crosstalk value for Atto643 into Cy3B.

**Supplementary figure 5:**
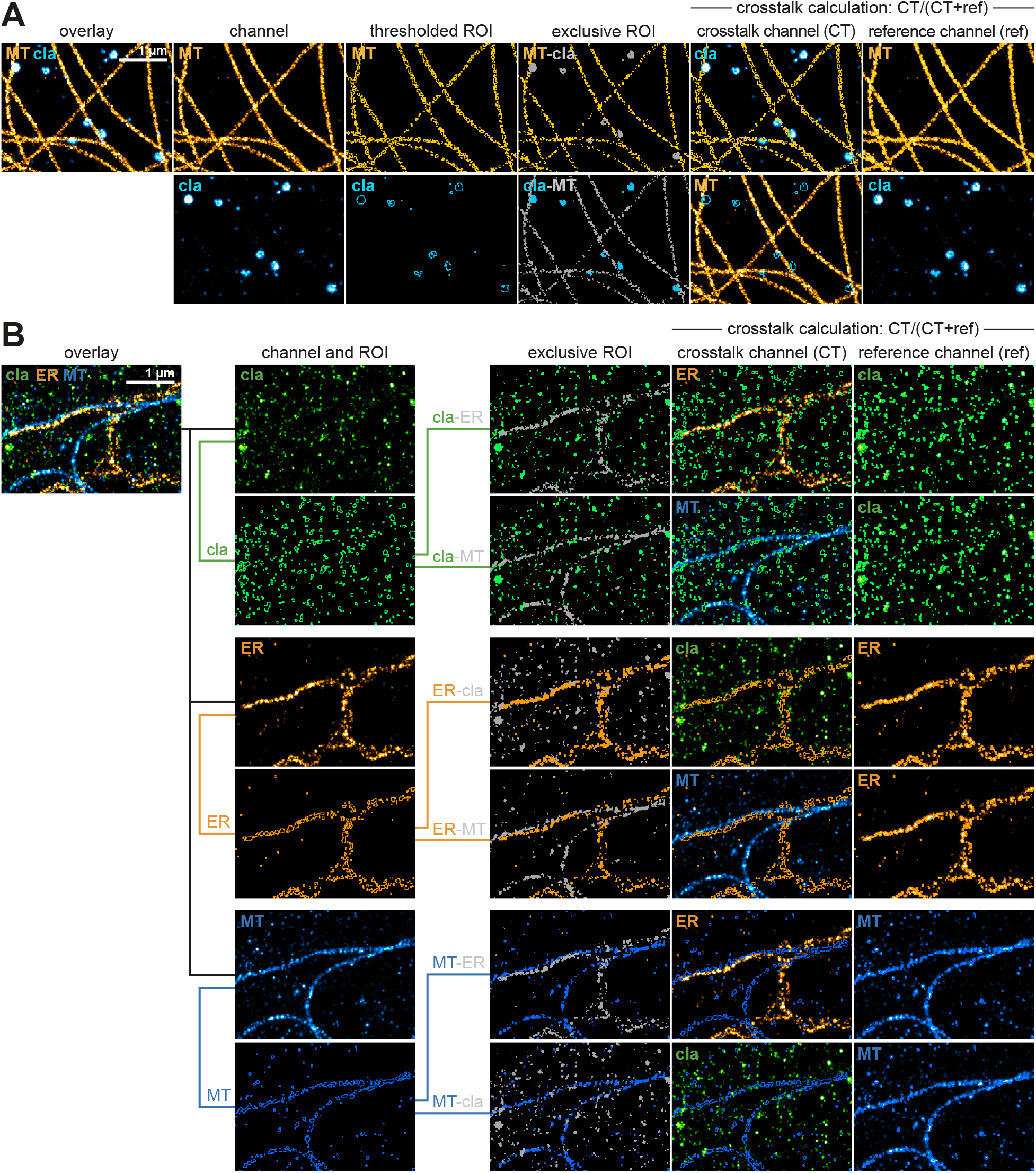
Generation of exclusive ROIs on multi-target images. **A.** Exclusive ROI crosstalk measurement procedure for a 2-color image of a COS cell stained for microtubules (orange) and clathrin (cyan). From each channel (second column), ROIs are defined by thresholding (third column). Subtraction of each ROI by the other results in exclusive ROIs for each channel (fourth column), that are applied to the crosstalk channel (CT, non-target, fifth column) and to the reference channel (target, sixth column) to calculate the crosstalk value from the number of localizations inside the exclusive ROI in each channel. **B.** Exclusive ROI crosstalk measurement procedure for a 3-color image of a COS cell stained for clathrin (green), ER (orange) and microtubules (cyan). From each channel (second column, top images), ROIs are defined by thresholding (second column, bottom images). Subtraction of each ROI by the others results in two exclusive ROIs for each channel (third column), that are applied to their respective crosstalk channel (CT, non-target, fourth column) and to the reference channel (target, fifth column) to calculate the crosstalk value from the number of localizations inside the exclusive ROI in each channel.

**Table S1:**
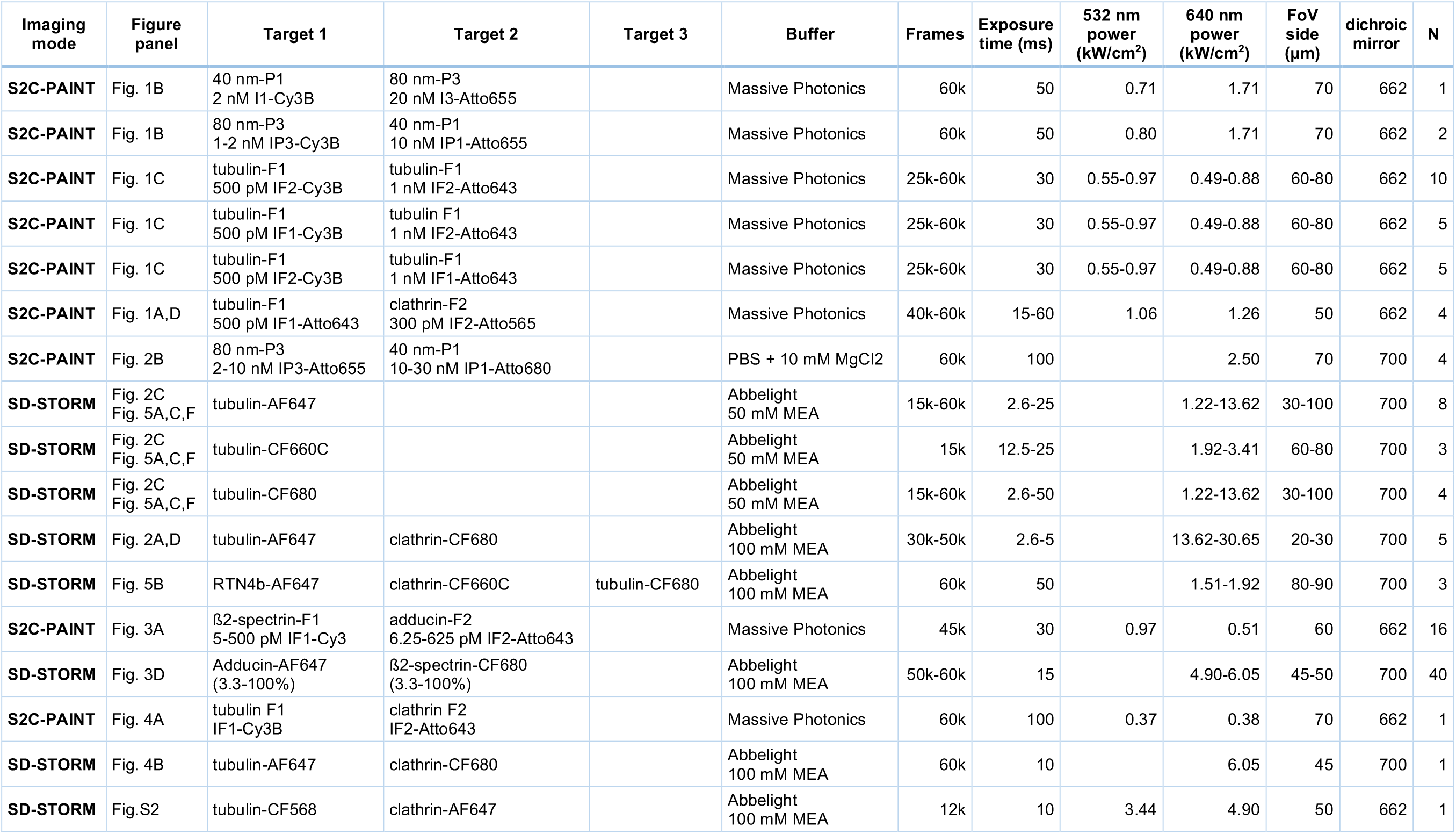
Acquisition parameters.

**Table S2:**
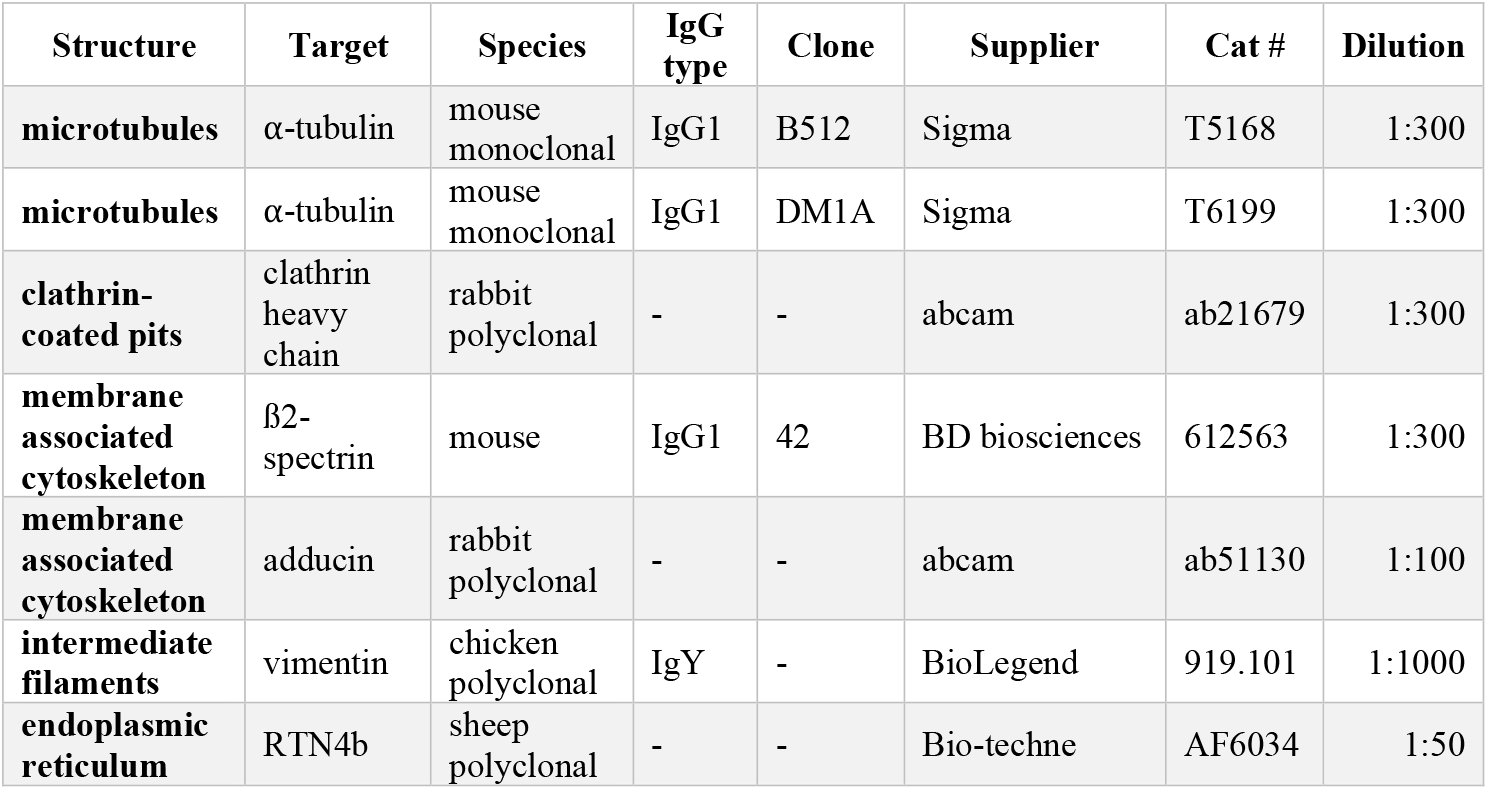
primary antibodies used for immunostaining.

**Table S3:** secondary antibodies used for immunostaining.

| Application | Host species | Target species | Conjugation | Supplier | Cat # | Dilution |
| --- | --- | --- | --- | --- | --- | --- |
| DNA-PAINT | donkey | mouse | F1 sequence | Massive Photonics | custom | 1:100 |
| DNA-PAINT | donkey | rabbit | F2 sequence | Massive Photonics | custom | 1:100 |
| STORM | donkey | mouse | Alexa Fluor 647 | ThermoFisher | A31571 | 1:300 |
| STORM | donkey | rabbit | Alexa Fluor 647 | Jackson ImmunoResearch | 711-605-152 | 1:300 |
| STORM | goat | chicken | Alexa Fluor 647 | Invitrogen | A21449 | 1:300 |
| STORM | donkey | mouse | CF660C | Biotium | BTM20815-500UL | 1:150 |
| STORM | donkey | rabbit | CF660C | Biotium | BTM20816-500UL | 1:150 |
| STORM | Donkey | Mouse | CF680 | Biotium | BTM20819-500UL | 1:150 |
| STORM | Donkey | Rabbit | CF680 | Biotium | BTM20820-500UL | 1:150 |

